# Faster detection of “darks” than “brights” by monkey superior colliculus neurons

**DOI:** 10.1101/2022.08.03.502615

**Authors:** Tatiana Malevich, Tong Zhang, Matthias P. Baumann, Amarender R. Bogadhi, Ziad M. Hafed

**Author notes:** Co-first authors (Tatiana Malevich, Tong Zhang, Matthias P. Baumann). Correspondence to: Tatiana Malevich and Ziad M. Hafed Werner Reichardt Center for Integrative Neuroscience Otfried-Müller Str. 25, Tübingen, Germany, 72076.

## Abstract

Visual processing is segregated into ON and OFF channels as early as in the retina, and the superficial (output) layers of the primary visual cortex are dominated by neurons preferring dark stimuli. However, it is not clear how the timing of neural processing differs between “darks” and “brights” in general, especially in light of psychophysical evidence; it is also equally not clear how subcortical visual pathways that are critical for active orienting represent stimuli of positive (luminance increments) and negative (luminance decrements) contrast polarity. Here, we recorded from all visually-responsive neuron types in the superior colliculus (SC) of two male rhesus macaque monkeys. We presented a disc (0.51 deg radius) within the response fields (RF’s) of neurons, and we varied, across trials, stimulus Weber contrast relative to a gray background. We also varied contrast polarity. There was a large diversity of preferences for darks and brights across the population. However, regardless of individual neural sensitivity, most neurons responded significantly earlier to dark than bright stimuli. This resulted in a dissociation between neural preference and visual response onset latency: a neuron could exhibit a weaker response to a dark stimulus than to a bright stimulus of the same contrast, but it would still have an earlier response to the dark stimulus. Our results highlight an additional candidate visual neural pathway for explaining behavioral differences between the processing of darks and brights, and they demonstrate the importance of temporal aspects in the visual neural code for orienting eye movements.

**Significance statement:** Objects in our environment, such as birds flying across a bright sky, often project shadows (or images darker than the surround) on our retina. We studied how primate superior colliculus (SC) neurons visually process such dark stimuli. We found that the overall population of SC neurons represented both dark and bright stimuli equally well, as evidenced by a relatively equal distribution of neurons that were either more or less sensitive to darks. However, independent of sensitivity, the great majority of neurons detected dark stimuli earlier than bright stimuli, evidenced by a smaller response latency for the dark stimuli. Thus, SC neural response latency can be dissociated from response sensitivity, and it favors the faster detection of dark image contrasts.

## Introduction

Early visual processing is segregated into parallel pathways conveying information about either luminance increments or decrements in visual scenes (Hartline, 1938; Schiller et al., 1986). Such segregation starts in the retina and persists in the early retino-geniculate visual pathway (Hubel and Wiesel, 1961; Schiller et al., 1986). Interestingly, such segregation is also accompanied by asymmetries with which dark and bright stimuli are processed. For example, primate retinal ganglion cells possess asymmetric spatial and temporal properties depending on whether they are part of the ON or OFF pathway (Chichilnisky and Kalmar, 2002). Similarly, in the primate lateral geniculate nucleus (LGN), neurons with OFF-center response fields (RF’s) are more sensitive to their preferred stimuli (dark contrasts) than neurons with ON-center RF’s experiencing bright contrasts (Jiang et al., 2015). OFF-center neurons also have higher spontaneous activity and more sustained responses during visual stimulation (Jiang et al., 2015). Ultimately, signals reach the primary visual cortex (V1), where ON/OFF asymmetries are amplified. For example, primate V1 is dominated by “black” responses, especially in the superficial cortico-cortical output layers (Yeh et al., 2009).

Asymmetries in the processing of dark versus bright stimuli might make ecological sense. For example, the incidence of dark contrasts in natural scenes is not necessarily uniform. Instead, there is a coincidence of dark contrasts with regions of low spatial frequency, high contrast, and far binocular disparities in natural images (Cooper and Norcia, 2015). As a result, rhesus macaque V1 neurons having far preferred binocular disparities tend to also prefer dark contrasts (Samonds et al., 2012). Similarly, in cat V1, there is a systematic contrast-dependent OFF-dominance, matching natural scene statistics (Liu and Yao, 2014), and cat V1 neurons are more strongly driven by luminance decrements than increments at low spatial frequencies (Kremkow et al., 2014). Interestingly, cat studies revealed that ON and OFF domains in the LGN also exist in the V1 projections (Jin et al., 2008), with area centralis representations being dominated by dark preferences. Moreover, OFF-dominated LGN (Jin et al., 2011) and V1 (Komban et al., 2014) neurons respond earlier than ON-dominated ones. These last observations on OFF and ON channel timing are consistent with a large body of psychophysical literature for better and faster processing of dark stimuli (e.g. Komban et al., 2011; Komban et al., 2014).

Having said that, whether monkey superior colliculus (SC) neurons differentially process dark stimuli remains unclear. In the mouse SC, the majority of superficial layer neurons prefer dark stimuli (De Franceschi and Solomon, 2018), consistent with the RF subfield structure of these neurons (Wang et al., 2010). Yet, it is not clear whether such dark preference still exists in the deeper SC layers, and whether it is accompanied by differences in visual response latencies. Moreover, differences in the ecological environments and neuroanatomical organizations of mice and other species do not trivially predict how primate SC neurons might behave with respect to luminance contrast polarity. Therefore, we exhaustively characterized all visually-responsive rhesus macaque monkey SC neurons (that is, also including intermediate and deeper layer neurons). We were particularly motivated by our recent observations of differential effects of contrast polarity on microsaccades (Malevich et al., 2021).

In contrast to LGN, V1, and SC results from other species, we did not find a dominant preference for dark stimuli in the primate SC. Rather, there was significant diversity, with approximately half of the neurons being more sensitive to bright stimuli. Moreover, at high contrasts, SC neurons tended to prefer bright rather than dark stimuli, with this trend disappearing for the lowest contrasts. Despite such diversity, what we did find was that the majority of SC neurons had significantly shorter visual response latencies to dark stimuli. Thus, there was a dissociation between visual response latency and visual response sensitivity, reminiscent of a similar dissociation that we observed in the case of spatial frequency tuning (Chen et al., 2018). Such a dissociation was sufficient to account for at least some saccadic reaction time dependencies on stimulus luminance polarity in our experiments.

## Materials and methods

### Experimental animals and ethics approvals

We recorded superior colliculus (SC) neural activity from two adult, male rhesus macaque monkeys (M and A) aged 9 and 10 years, and weighing 9.5 kg and 10 kg, respectively. We also measured saccadic reaction times from the same two animals plus a third one (F; aged 11 years and weighing 14 kg). The experiments were approved by ethics committees at the regional governmental offices of the city of Tübingen.

### Laboratory setup and animal preparation

The experiments were conducted in the same laboratory as that described in our recent studies (Bogadhi et al., 2020; Bogadhi and Hafed, 2022). Briefly, the monkeys were seated in a darkened booth approximately 72 cm from a calibrated and linearized CRT display spanning approximately 31 deg horizontally and 23 deg vertically. For monkey F only, the display was an LCD device running at 138 Hz (AOC AG273QX2700, 27”), as in (Malevich et al., 2021). Data acquisition and stimulus control were managed by a custom-made system based on PLDAPS (Eastman and Huk, 2012). The system integrated a DataPixx display control device (VPixx Technologies, Inc.) with the Psychophysics Toolbox (Brainard, 1997; Pelli, 1997; Kleiner et al., 2007) and an OmniPlex neural data processor (Plexon, Inc.).

The monkeys were prepared for behavioral training and electrophysiological recordings earlier (Tian et al., 2018; Buonocore et al., 2019; Skinner et al., 2019; Malevich et al., 2020). Specifically, each monkey was implanted with a head-holder, and monkeys M and A were also implanted with a scleral search coil in one eye. The search coil allowed tracking eye movements using the magnetic induction technique (Fuchs and Robinson, 1966; Judge et al., 1980), and the head-holder comfortably stabilized head position during the experiments. Eye movements in monkey F were recorded with a video-based eye tracker (EyeLink1000; desktop mount; 1 KHz sampling rate). For the present experiments, monkeys M and A also each had a recording chamber centered on the midline and tilted 38 deg posterior of vertical, allowing access to both the right and left SC (Bogadhi and Hafed, 2022).

### Behavioral tasks

For the recording data in monkeys M and A, we employed a gaze fixation task in which we presented static disc of 0.51 deg radius within the visual response field (RF) of a recorded neuron. Each trial started with the onset of a black (0.11 cd/m^2^) fixation spot at screen center. After 550-800 ms of stable fixation on the spot, the disc appeared and remained on for at least ~500 ms. In each trial, the disc could have a Weber contrast of 5%, 10%, 20%, 50%, or 100%. We defined Weber contrast as |*I_s_-I_b_*|/*I_b_*, where *I_s_* is the disc’s luminance value and *I_b_* is the gray background’s luminance value. We often described the contrast as a percentage for convenience (e.g. 5% contrast). Importantly, across trials, the disc could have either positive or negative luminance polarity relative to the gray background, meaning that *I_s_* could be either higher (positive polarity) or lower (negative polarity) than *I_b_*. The gray background had a luminance (*I_b_*) of 25.09 cd/m^2^. We collected approximately 50 trials per condition per neuron.

For some neurons in both monkeys (sometimes in the very same sessions as in the above task), we also ran an immediate orienting version of the stimulus polarity task. That is, at the time of disc onset, we extinguished the fixation spot (which was now white instead of black) simultaneously. This instructed the monkeys to generate an immediate orienting saccade towards the disc. We used this task to confirm that initial visual responses in the main task above were not dictated by the black fixation spot at display center, since the current task had a white fixation spot and showed similar observations (see Results), and also to obtain saccadic reaction time data for additional behavioral analyses (see below). Also note that, for neurophysiological analysis purposes, we only analyzed the initial visual response in this task. Saccade-related responses were deferred to another unrelated project focusing on SC motor bursts, and they are not described here. Finally, to reduce trial counts in this task, we only tested three contrast levels (10%, 50%, and 100%). We collected approximately 50 trials per condition per neuron.

For exploring a potential behavioral consequence of faster detection of dark stimuli by SC neurons (which we describe in Results), we tested our three monkeys on the saccadic reaction time version of the task, which we just described above. For monkeys M and A, we analyzed reaction times from the same sessions as those collected during neurophysiological recordings. Stimulus locations were, thus, dictated by recorded neurons’ response field (RF) locations. For monkey F, we ran behavior-only sessions. In this case, we randomly varied stimulus locations across 4 diagonals, with an eccentricity of 8.9 deg. We analyzed a total of 457-773 saccades per condition per monkey for our behavioral reaction time analyses.

### Neurophysiological procedures

For most experiments, we recorded SC neurons using linear electrode arrays inserted across the SC depth (24 channel V Probes; Plexon, Inc.). For some experiments, we also used single tungsten electrodes, in which we targeted and isolated individual neurons online during the experiments. In all cases, including the single electrode sessions, we also performed offline sorting to re-isolate neurons for inclusion in the data analysis pipeline (Pachitariu et al., 2016). Sorting and general data analysis pipeline details were similar to those described recently (Bogadhi and Hafed, 2022).

Before collecting data from our main tasks, we first identified the SC by running RF mapping tasks. These included delayed and memory-guided saccades (Chen et al., 2015; Chen and Hafed, 2017). The mapping tasks allowed us to select the stimulus location for our main experiments, and also to confirm that our neurons possessed visual responses (or stimulus-triggered inhibition). For the simultaneous recordings of multiple neurons with electrode arrays, we picked a disc location that we felt lay within the RF’s of most neurons that we could identify online. This was possible given that our electrodes were penetrating the SC surface at a quasi-orthogonal angle, meaning that the RF’s at different depths generally had similar locations. Also note that for all analyses, we were always interested in comparing responses to bright and dark discs at the very same location. That is, our comparison of interest was the luminance polarity at a given RF location for a given neuron. In separate experiments, we mapped RF’s with positive and negative luminance polarity spots, but these data will be described in detail separately. For the present purposes, suffice it to say that all RF’s had sensitivity to both black and white targets at their center, justifying our current comparison of response sensitivity at a single given RF location per neuron.

Across all experiments, we recorded from neurons with extrafoveal eccentricities (e.g. 2.1-20 deg preferred eccentricity across the population), meaning that we often presented stimuli far from the fixation spot.

### Eye movement data analysis

We detected saccades and microsaccades as described previously (Chen and Hafed, 2013; Bellet et al., 2019). We used the detections for two primary purposes. First, in the recording tasks, we excluded all trials in which there were microsaccades occurring within an interval from −50 ms to +50 ms relative to stimulus onset. This allowed us to measure baseline visual responses that were not modulated by the known influences of microsaccades on SC activity (Hafed and Krauzlis, 2010; Chen et al., 2015; Chen and Hafed, 2017). We also performed the same filtering for behavioral analyses of saccadic reaction times. Second, for the behavioral analyses, we used saccade detection to measure saccadic reaction times towards the dark and bright stimuli.

To identify a response saccade and subsequently analyze its reaction time, we required that it had a latency of 50 to 500 ms from stimulus onset, and that it was directed towards the stimulus (this latter criterion was easy to achieve because we used computer-controlled reward windows around the target to allow rewarding the monkeys based on successful saccade generation towards the target). In all neural and behavioral analyses, we also excluded trials with blinks or other movement artifacts near stimulus onset. Statistically, we were interested in whether contrast or luminance polarity affected saccadic reaction times. Therefore, we performed a 1-way non-parametric ANOVA (Kruskal-Wallis test) in each monkey testing for the effect of contrast (d.f.: 2), when collapsing across luminance polarities, on the monkey’s reaction times. Similarly, we also performed a Kruskal-Wallis test exploring the effect of luminance polarity (d.f.: 1), when collapsing across contrasts, on reaction times.

### Neural data analysis

We analyzed a total of 221 SC neurons (109 from monkey M and 112 from monkey A) from the fixation task. We also analyzed 225 neurons from the immediate saccade version of the task (113 from monkey M and 112 from monkey A). Ninety of the neurons in the second task (all in monkey A) were also recorded from the fixation variant of the task.

The bulk of our analyses was on neurons exhibiting a positive visual response to stimulus onset (that is, an increase in firing rate relative to baseline shortly after stimulus onset). We, therefore, first tested for the presence of a positive visual response (or burst) after stimulus appearance. In each neuron, we defined a baseline interval as the final 50 ms before stimulus onset. We then defined a visual response interval as the time interval 10-200 ms after stimulus onset. Across all repetitions of a given stimulus condition (e.g. 100% contrast; positive polarity), we measured average firing rate in the response interval and statistically compared it to average firing rate in the baseline interval. If the response interval firing rate, across repetitions of a given condition, was statistically significantly larger (one-tailed, paired t-test; p<0.025) than baseline firing rate, and if this significance occurred for both polarity conditions (dark and bright) and with absolute Weber contrasts of 50% and 100%, then we considered the neuron to have a positive visual response to stimulus onset. We did not include lower contrast trials (whether positive or negative polarity) in assessing for the presence of visual responses because some neurons, even when having strong visual bursts for high contrast stimuli, did not respond to such lower contrasts. Across our population in the fixation task, we had a total of 172 neurons (92 from monkey M and 80 from monkey A) exhibiting visual bursts after stimulus onset with the above criteria. For the immediate saccade version of the task, we only analyzed neurons with a positive visual burst in the interval 10-180 ms after stimulus onset; this resulted in a total of 213 neurons (109 from monkey M and 104 from monkey A).

For a subset of neurons, stimulus onset caused a transient decrease in firing rate from baseline, rather than an increase. We performed analyses of these neurons as well, from the fixation task only. To assess the neurons as having a transient decrease in firing rate that was time-locked to stimulus onset, we repeated the same procedure above, but we now checked for a statistically significant decrease in firing rate in the response interval, rather than an increase. We analyzed 15 neurons with transiently decreasing firing rates immediately after stimulus onset (9 from monkey M and 6 from monkey A).

To obtain contrast sensitivity curves from the neurons with visual bursts, we measured the peak value of the average firing rate curve in a response interval after stimulus onset. Since visual response latency in the SC varies with stimulus contrast (Li and Basso, 2008; Marino et al., 2012; Marino et al., 2015), we tailored the measurement interval for each contrast as follows: 15-105 ms after stimulus onset for 100% contrast; 20-110 ms after stimulus onset for 50% contrast; 20-115 ms after stimulus onset for 20% contrast; 35-125 ms after stimulus onset for 10% contrast; and 45-135 ms after stimulus onset for 5% contrast. Note that we used the same measurement intervals for all neurons and also for both positive and negative polarity stimuli. Even though we found a difference in response latency between positive and negative polarity stimuli (as described in Results below), our measurement intervals were large enough to encompass (and exceed) any such latency differences. Therefore, our estimates of contrast sensitivity for brights and darks were not biased by using similar measurement intervals for both types of stimuli (especially because we were searching for only the peak firing rate). After measuring firing rates in the above intervals for each contrast, we plotted the measured firing rates as a function of absolute contrast. We then fit contrast sensitivity curves using the following equation:

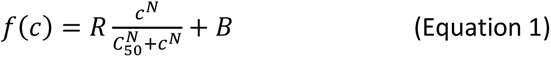

where *f* is the estimated firing rate, *c* is stimulus contrast, *C50* is semi-saturation contrast, *R* is the dynamic range of the response, *N* is the sensitivity/slope of the contrast sensitivity curve, and *B* is the baseline firing rate (which we just measured across all trials from the same baseline interval mentioned above; final 50 ms before stimulus onset). We then compared the fit parameters *R*, *C50*, and *N* for either bright or dark stimuli to assess whether there were differences in contrast sensitivity between them in the SC. We did this by computing parameter modulation indices as a function of luminance contrast polarity. For example, to compare how *R* was modulated by luminance polarity, we calculated the *R* parameter for bright stimuli minus the *R* parameter for dark stimuli, and we divided this difference by the sum of *R* values for bright and dark stimuli. This gave us a value between −1 and +1. We then plotted histograms of parameter modulation indices across the population.

For assessing the time course of changes of contrast sensitivity in the sustained interval long after stimulus onset (that is, after the initial visual burst), we again obtained fits of equation 1 but now based on measurements after the initial visual burst. To do so, we exploited the fact that our firing rate estimates were already averaging across time (because of a convolution of spike times with a gaussian of σ 40 ms). Therefore, for each time after 80 ms after stimulus onset and before 220 ms, we used instantaneous average firing rate as a measure to input to the fit of equation 1 for a given contrast. This allowed us to obtain time courses of semi-saturation contrast (*C50*), sensitivity/slope (*N*), and dynamic range (*R*) during the sustained interval long after stimulus onset. Note that even though the intervals that we chose for the sustained response analysis slightly overlapped with the intervals that we picked up for the initial visual burst analyses mentioned above, the latter analyses were performed on the peak values of the average firing rates, which definitely belonged to the earlier phases of the neural responses (and typically occurred earlier than 80 ms); that is, the initial visual burst intervals were just ranges meant to catch the peak response. Also note that the above contrast sensitivity fits were only performed on the fixation version of the task because we could obtain a longer period of sustained response than in the immediate saccade version of the task.

For estimating visual response latency in both tasks, we measured the firing rate of a given neuron in a baseline interval (final 50 ms before stimulus onset) across all trials. Then, for each condition (e.g. bright luminance polarity; 20% contrast), we marched forward in time after stimulus onset until 300 ms (typically, the algorithm converged on a visual burst much earlier, of course). As soon as the lower bound of the 95% confidence interval around the average firing rate of the neuron across trial repetitions exceeded the average baseline activity (and continued to do so for at least 30 ms), we flagged the time as the response latency of the neuron. Whenever this algorithm failed to detect the response latency in at least one of the luminance polarities for a given stimulus contrast level, we excluded the neuron from further analysis in that contrast level. This explains the varying numbers of neurons reported in some figures (e.g. the different panels of Fig. 4 in Results). We then compared such response latency across contrasts and stimulus polarities. Note that we focused on relative latency differences across luminance polarities in our analyses. This is important to note because firing rate estimates (in our case, convolution of spike times with a Gaussian kernel) necessarily blurs the exact response onset times of the neurons. However, our approach of estimating response latencies described above still captured the latency differences that we were interested in documenting, and it simplified the detection of visual response latencies for neurons with non-zero baseline firing rates.

To statistically test for differences in latencies between luminance polarities at a given contrast level, we used non-parametric permutation tests on the pairwise mean latency differences, with 10000 permutations. That is, we obtained the permutation distribution by shuffling the polarity labels of the latencies for 10000 times while maintaining their pairwise relationship and calculating their pairwise difference. Monte Carlo p-values were obtained by assessing the probability of getting larger than or equal to absolute latency differences in the permutation distribution than the absolute latency difference of the original data. We ran the tests separately for each monkey to ensure that our pooling of data in figures for visualization purposes was justified.

We also used a similar approach to test for statistically significant effects of upper versus lower visual field RF location on the latency differences between luminance polarities. This time, we obtained the latency differences between the responses to bright and dark stimuli, and then subtracted this measurement for the upper visual field neurons from the measurement for the lower visual field neurons. We defined lower and upper visual field neurons based on the location of the stimulus (which was placed close to the location of the RF hotspot location). Thus, negative values in the final measurement would indicate a larger difference between dark and bright stimuli in the upper visual field than in the lower visual field. After that, we ran permutation tests by shuffling the labels of the upper and lower visual field neurons for 10000 times. To assess the significance of the results, we calculated the Monte Carlo p-value. The same procedure was applied to assess the absolute values of sensitivity differences (see next paragraph) between dark and bright stimuli in the upper and lower visual fields.

To compare visual response latency to sensitivity, we also measured peak firing rate in the initial visual response interval (as defined for each contrast above) of the neuron. First, to test whether there was an effect of luminance polarity on sensitivity, we used permutation tests in the same way as we did for the latency analysis described above, but this time on the pairwise mean peak response differences, separately for each monkey and contrast level. We then checked whether there was a dissociation between response latency and sensitivity (i.e. response strength) for black and white stimuli, as we previously saw for spatial frequency stimuli (Chen et al., 2018). We did so by sorting the neurons according to the difference in response latencies between brights and darks, and then checking whether the same sorting applied to the difference in response sensitivities. Further, we pooled the data across monkeys and calculated Pearson’s correlation coefficients between differences in peak visual responses and differences in visual response latencies, separately for each of the contrast levels (see Results).

For the immediate saccade version of the task, we only analyzed initial visual bursts (50-130 ms after stimulus onset) and not sustained intervals. This was because the response saccade occurred too soon after the initial visual bursts. We assessed both response sensitivity and response latency (as described above) to confirm that we got similar results to those from the fixation task.

For the neurons with transient decreases in firing rate, we assessed response latency in a similar way to the neurons with visual bursts, but we looked for statistically significant decreases in firing rate after stimulus onset, rather than increases.

In some figures, for illustration and visualization purposes, we elected to show example population firing rates from individual monkeys. For example, we did this in Fig. 8A, B in Results. To obtain such population firing rates, we obtained the normalized average firing rate of each neuron, per monkey and condition. That is, for each neuron, we found the peak visual response in the interval 0-100 ms after stimulus onset for the 100% contrast stimuli, regardless of the stimulus polarity. Then, for each contrast and polarity, we normalized the neuron’s average firing rate by that peak visual response value. This resulted in a series of average normalized curves for the neuron across conditions. After that, we averaged all of the normalized firing rate curves of each monkey’s neurons in a given condition. This gave us a population summary of responses, maintaining the relative changes in responses across conditions. We used a similar approach in Fig. 11B, D.

### Experimental design and statistical analyses

We recorded neurons in an unbiased manner by collecting data in parallel (with linear electrode arrays) in most sessions and then sorting the neurons offline. This allowed us to minimize sampling bias. In each variant of the task, we also analyzed >80 neurons per monkey. This provided a large enough sample to assess the reliability of our interpretations. Within each neuron, we ensured collecting approximately 35-50 repetitions per condition (after filtering out bad trials and so on) to allow robust within-neuron statistics. Similarly, in our behavioral analyses, we collected thousands of saccades. In all cases, we randomly interleaved stimulus presentations across trials, to avoid any blocking effects.

We provided descriptive statistics in all figures, showing numbers of observations and measures of variability. Also, in most of our critical analyses (e.g. Figs. 4–6 in Results), we showed the full distributions of data points that we had.

Since the replicate of interest was neurons, our numbers of sampled neurons were sufficient. The use of two monkeys in recording was valuable to increase neuron counts, and to also demonstrate repeatability across individuals. Our results were highly similar in the two animals (e.g. Fig. 8A, B in Results). When they did differ, we showed each individual monkey’s results separately (e.g. Fig. 11 in Results), and this was highly useful for us to interpret the behavioral results. Moreover, we collected behavior from a third monkey exactly to improve our interpretation of our individual monkey behavioral phenomena.

All statistical tests are reported and justified in Results at appropriate points in the text. As stated above, we statistically analyzed each monkey’s data individually, confirming that each monkey showed the same effects (unless otherwise stated; for example, in Fig. 11 in Results).

## Results

We investigated how monkey superior colliculus (SC) neurons respond to dark and bright visual stimuli. In our primary task, the monkeys fixated while we presented a small disc that was either higher or lower in luminance than the gray background of the display. We varied the contrast of the disc from the background luminance, and we assessed contrast sensitivity curves separately for positive and negative luminance polarities. We first analyzed the neurons that exhibited visual bursts (that is, increases in firing rate) after stimulus onset, and we investigated visual burst strength, visual burst latency, as well as sustained response dynamics for dark and bright stimuli. The results for these neurons are described next, followed by an analysis of a smaller number of neurons for which stimulus onsets caused transient decreases in firing rates, rather than increases.

### Diverse preferences for darks and brights across SC neurons

We first asked whether neurons tended to be more sensitive to darks or brights across the population. For each recorded neuron, we plotted firing rate as a function of time from stimulus onset, and we assessed the strength of the visual burst as a function of luminance contrast polarity. Figure 1A-C shows the responses of three example neurons (from the same monkey, A) to a 100% contrast stimulus. The black lines show responses to the negative polarity stimulus (darker than background), and the light gray lines show responses to the positive polarity stimulus (brighter than background). In all cases, the negative and positive polarity stimuli were of the same size and presented at the same location. They also had the same absolute Weber contrast, and their presentation sequence was randomly counterbalanced across trials. As can be seen, there was a diversity of neural preferences across the three neurons: neuron 1 (Fig. 1A) was more sensitive to the positive polarity stimulus than to the negative polarity stimulus; neuron 2 was, more or less, equally sensitive to the two stimuli (Fig. 1B); and neuron 3 was clearly more sensitive to the dark stimulus (Fig. 1C). We also plotted full contrast sensitivity curves for the same neurons (Fig. 1D-F) by relating peak visual response strength to stimulus contrast (Methods). Consistent with Fig. 1A-C, there was a diversity of preferences for darks and brights across the three neurons in their full contrast sensitivity curves.

**Figure 1.**
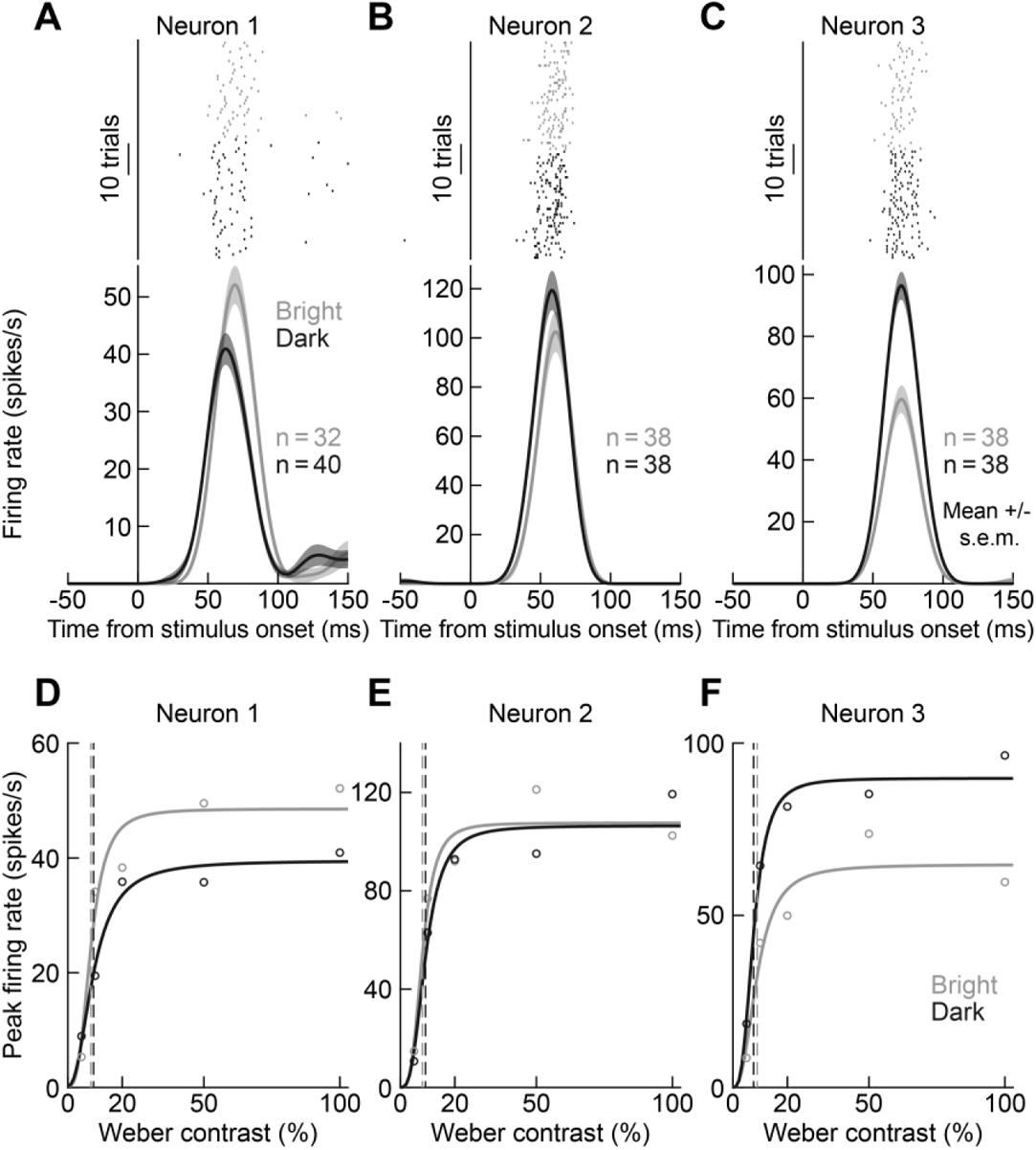
Diversity of preferences, in terms of visual burst strength, for dark and bright stimuli by SC neurons. **(A-C)** Visual responses of three example neurons from monkey A to a 100% contrast stimulus appearing within their response fields (RF’s). Black indicates that the stimulus was negative in luminance polarity (darker than the background); gray indicates that the stimulus was brighter than the background. Each row of tick marks indicates a single trial, and each tick mark indicates the time of an action potential. The firing rate plots below the raster plots summarize the neurons’ firing rates. Neuron 1 had a higher sensitivity to bright stimuli, whereas neuron 2 was equally sensitive to dark and bright stimuli. Neuron 3, on the other hand, clearly preferred dark stimuli. Note that response latency (that is, when the stimulus-evoked action potentials first appeared) was always shorter for dark stimuli (see subsequent analyses). Error bars denote s.e.m. across trials. **(D-F)** For each neuron, we measured peak average firing rate after stimulus onset (individual symbols), and we plotted it as a function of stimulus Weber contrast. We also fit the data with continuous curves (Methods). Neuron 1 had higher contrast sensitivity for bright stimuli, evidenced by the higher plateau firing rate at maximal contrast. Neuron 2 plateaued at the same firing rate for both dark and bright stimuli, and neuron 3 was more sensitive to dark stimuli. Dashed vertical lines indicate *C50*, the semi-saturation contrast of each neuron (Methods).

These observations held across the population of 172 neurons that we analyzed. For each neuron, we fit a contrast sensitivity function (equation 1; see Fig. 1D-F for examples) by optimizing three parameters characterizing how the neuron altered its visual response with Weber contrast: *R* reflected the dynamic range of the response, *C50* characterized the semi-saturation contrast of the neuron, and *N* characterized the steepness of the contrast sensitivity curve (slope parameter). We performed such a fit for either positive or negative luminance polarity stimuli. We then obtained a parameter modulation index, describing, for each neuron, to what extent each parameter of the fit was different between positive and negative luminance polarity stimuli. For example, for dynamic range (parameter *R* in equation 1), we obtained the *R* value for bright stimuli minus the *R* value for dark stimuli in each neuron, and we then divided this difference by the sum of *R* values for the two stimulus types (Methods). This gave us an index in which 1 meant that the neuron responded maximally only to bright stimuli and −1 meant that the neuron responded maximally only to dark stimuli. An *R* parameter modulation index value of 0, instead, indicated equal visual response dynamic ranges for bright and dark stimuli. We then plotted histograms of the parameter modulation indices across the population. As can be seen in Fig. 2, all three modulation indices of the contrast sensitivity function fits had distributions straddling 0, and with large diversity across the population. Some neurons clearly preferred bright stimuli, others clearly preferred dark stimuli, and yet others were equally sensitive to darks and brights (near 0 in the histograms of Fig. 2). The vertical lines in Fig. 2 indicate the mean (solid) and median (dashed) parameter modulation index values across neurons, and they were all close to 0. Approximately half of the neurons were more sensitive to bright stimuli (whether in terms of dynamic range, semi-saturation contrast, or slope of the contrast sensitivity function), and the other half were more sensitive to dark stimuli.

**Figure 2.**
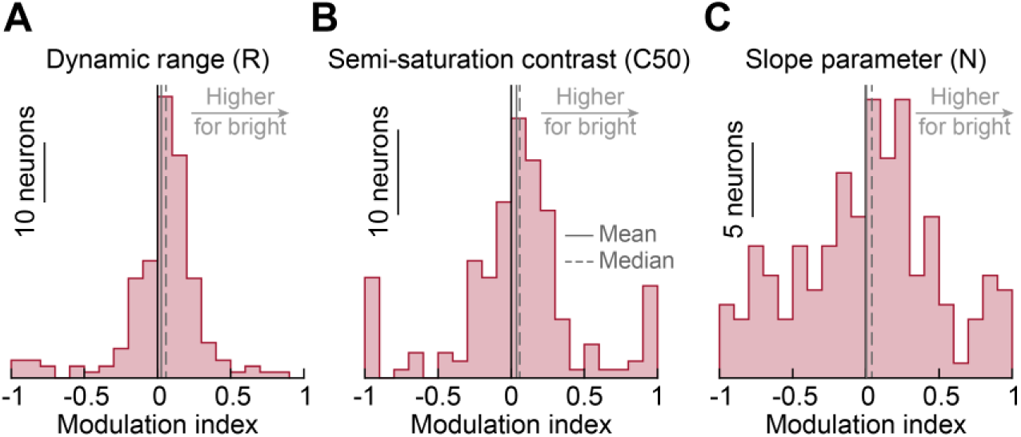
Diversity of contrast sensitivity curve parameters for dark and bright stimuli across SC neurons. **(A)** For each neuron, we compared the neuron’s visual response dynamic range (parameter *R* in equation 1; Methods) between dark and bright stimuli. A modulation index value of 1 indicates maximal responsiveness to only bright stimuli, and a value of −1 indicates maximal responsiveness only to dark stimuli. Most neurons responded to both stimuli (note the rarity of +/-1 modulation index values), but with varying degrees of sensitivity; some neurons clearly preferred dark stimuli, whereas others clearly preferred bright stimuli (similar to the examples in Fig. 1). The vertical lines show mean (solid) and median (dashed) modulation index values across the population, and they were both close to 0: across the population, SC visual responses were equally sensitive to dark and bright stimuli. **(B)** Similar observations for the semi-saturation contrast (*C50*) parameter. Neurons with positive modulation indices in this case are neurons with higher semi-saturation contrasts for bright stimuli (that is, they were less sensitive to bright than dark stimuli). Again, the population average and median semi-saturation contrasts were largely similar between darks and brights (vertical lines), but with large variability across individual neurons. **(C)** Similar observations for the slope parameter of the contrast sensitivity curves. These results are consistent with the example neurons of Fig. 1.

Therefore, in the SC, we noticed a substantial diversity of sensitivity preferences for darks and brights across the population (unlike in LGN and V1). This suggests that stimuli of both positive and negative luminance polarities can indeed be represented well by SC neural populations.

### Earlier detection of darks by SC neurons, regardless of preference

Unlike response sensitivity, for which we saw diverse preferences for brights and darks (Figs. 1, 2), SC neurons exhibited systematically shorter visual response latencies for dark stimuli, independently of their visual response strengths at a given contrast. Consider, for example, the same three neurons of Fig. 1A-C. In each of them, visual responses occurred earlier for the dark stimuli than for the bright stimuli, as can be visually assessed from the spike rasters and the firing rate density plots below them. This happened even for neuron 1, which preferred bright stimuli (Fig. 1A). It also happened at different contrast levels (Fig. 3), even though stimulus contrast expectedly modulated the response strength and latency of each neuron. For example, at 20% contrast, all three neurons from Fig. 1 still responded earlier to dark than bright stimuli, despite the weakened and delayed visual responses relative to the 100% contrast conditions. Thus, at each contrast level, there was an apparent dissociation between visual response sensitivity and visual response latency in these three neurons, not unlike what we recently observed when we presented different spatial frequencies to SC neurons (Chen et al., 2018).

**Figure 3.**
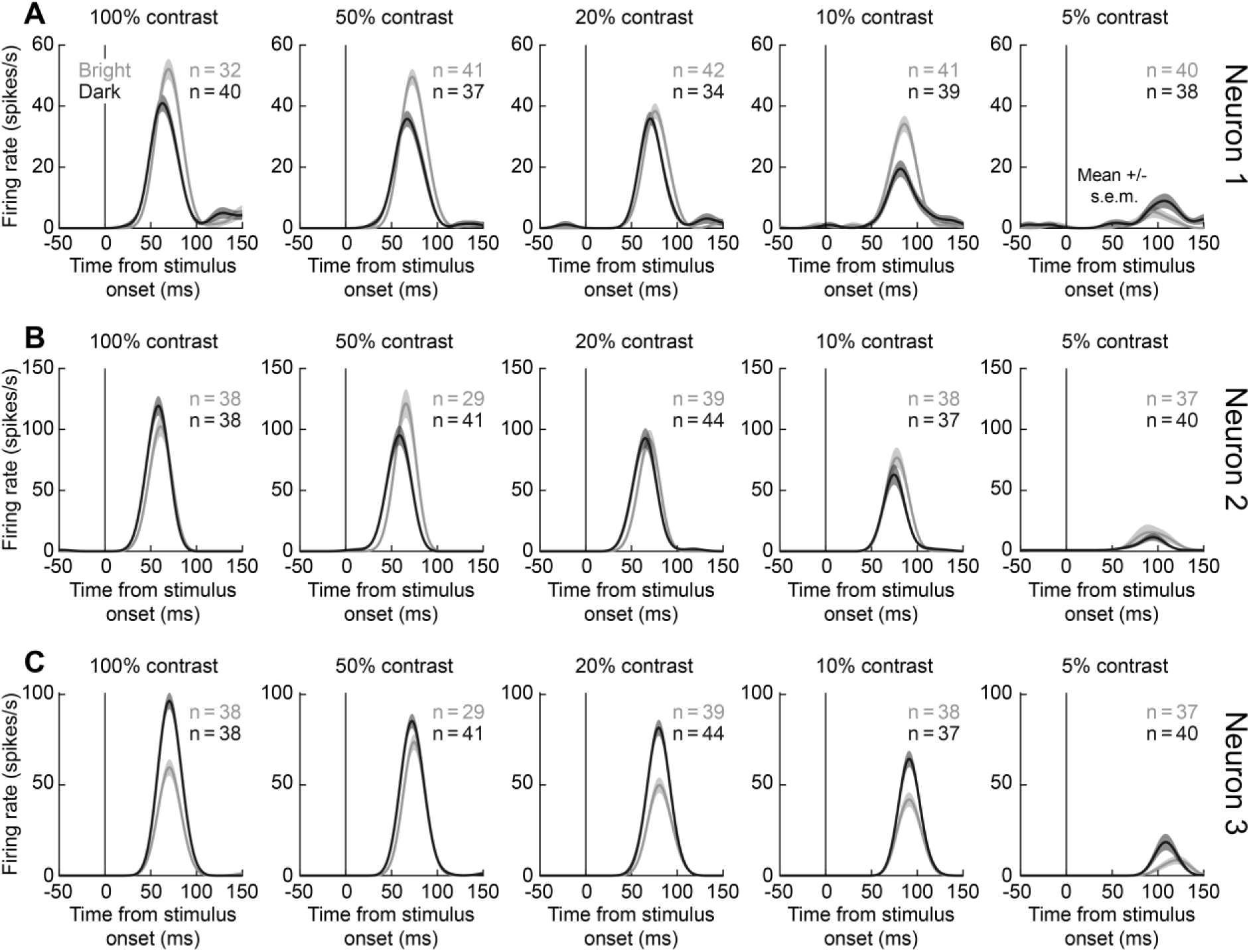
Earlier visual bursts for dark than bright stimuli at different stimulus contrasts. **(A-C)** Visual responses of the three example neurons of Fig. 1 at each of our tested contrasts (different columns). Black curves indicate dark stimuli, and gray curves indicate bright stimuli. Reducing contrast expectedly weakened and delayed visual responses for both dark and bright stimuli (compare firing rates across columns; also see Fig. 1D-F). Note, however, that at each contrast level, visual bursts occurred slightly earlier for dark than bright stimuli, even for neuron 1, which was more sensitive to bright stimuli. Also note that at the weakest contrast level (5%), both neuron 1 (**A**) and neuron 3 (**C**) were more sensitive to light decrements than light increments. We quantify these observations in subsequent figures and analyses. Error bars denote s.e.m. across trials.

To investigate this dissociation further, we estimated, for each neuron, the onset of the visual burst as the first time point at which the lower bound of the 95% confidence interval of the neuron’s average firing rate was elevated for a prolonged period of time above its baseline activity (Methods). Even though estimating neural response latencies from firing rate measures like we did might blur the actual absolute values of the response latencies, due to convolution kernels with spike times, this approach was still sufficient to capture the latency differences across luminance polarities that we were interested in (Methods). Therefore, for each contrast, we subtracted each neuron’s visual response latency for dark stimuli from its visual response latency for bright stimuli, and we sorted the neurons according to this difference. An example of such sorting can be seen in the top panel of Fig. 4A for the 100% contrast stimuli. Note how a majority of neurons (76.7%; 128 out of 167) had an earlier visual response for dark stimuli (evidenced by a positive latency difference in the figure). This is in contrast to the diversity of preferences for darks and brights seen in Figs. 1–3. In fact, with the very same sorting of the neurons as in the top panel, we next plotted (bottom panel of Fig. 4A) the same neurons’ differences in peak visual burst strengths between darks and brights (Methods). The neurons were no longer as properly ordered as in the top panel, suggesting that the latency effect in the top panel was not trivially explained by a systematic difference in response sensitivity between darks and brights. For example, both neurons 126 and 156 (highlighted in Fig. 4A with small diagonal arrows) possessed clearly stronger responses for bright stimuli than dark stimuli (positive difference in the bottom panel), but they both had a later response latency for bright stimuli (positive difference in the top panel). Therefore, visual responses to dark stimuli still occurred earlier than visual responses to bright stimuli even when neurons preferred bright stimuli.

**Figure 4.**
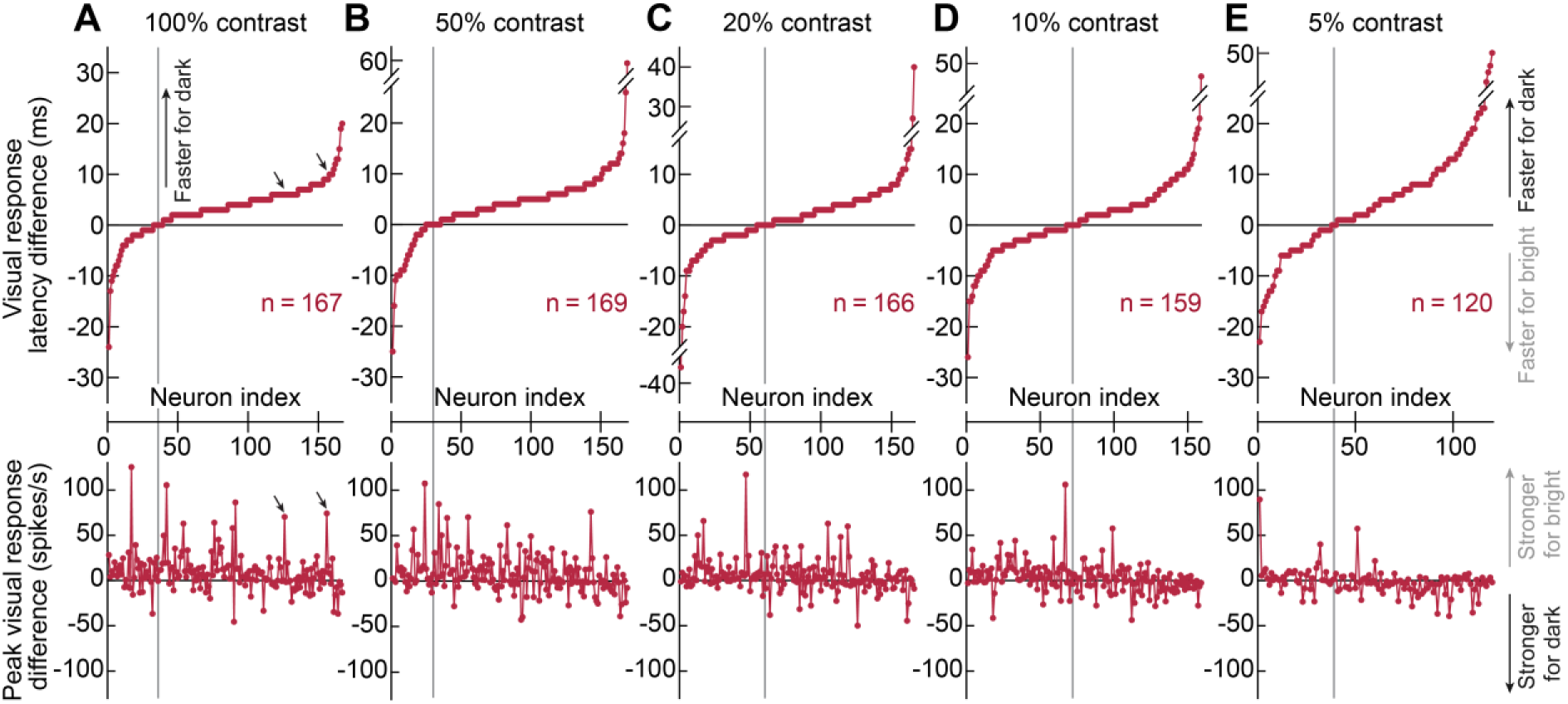
Faster detection of darks than brights by monkey superior colliculus neurons. **(A)** For the highest stimulus contrast, we estimated visual response latency (Methods) separately for dark and bright stimuli in each neuron. We then subtracted, for each neuron, the visual response latency for dark stimuli from the visual response latency for bright stimuli (positive would indicate earlier responses for dark stimuli). We then sorted all neurons based on this difference (top panel). In the bottom panel, we used the very same sorting, but we now plotted the difference in peak visual response strength between bright and dark stimuli (Methods). Most neurons had a shorter visual response latency for dark than bright stimuli (top panel; vertical line shows the sorted neuron index at which response latency differences flipped sign from negative to positive). This happened independently of response strength; the bottom panel (with the same sorting) did not show a systematic ordering. For example, the neurons highlighted with diagonal arrows preferred bright stimuli (bottom panel) but still detected dark stimuli earlier (top panel). **(B-E)** Similar results for lower contrasts. Of course, with lower and lower contrasts, the earlier detection of darks was less and less prevalent (see the crossover points in the top row). However, this was because lower contrasts were already associated with delayed and weakened visual responses (see Fig. 5). Also note that the lower row shows a decreasing likelihood of bright-preferring neurons as contrast level decreases, suggesting a contrast-dependent processing of darks and brights in SC neurons (see Fig. 6).

We also made similar observations for the other stimulus contrasts that we tested (Fig. 4B-E). Note that for each panel in Fig. 4, we indicated the total number of neurons included into each analysis, which varied across panels (that is, across contrast levels). This happened because some neurons may not have met our inclusion criteria for estimating visual response latencies, resulting in slightly different neuron counts across the different panels (Methods). For example, for the particularly low contrast stimuli (e.g. 5% and 10%), some neurons did not exhibit any significant visual bursts at all (Methods), so they were not included in the figure. Having said that, in all contrasts, there was a majority of neurons responding earlier to dark than bright stimuli (top row in each panel of Fig. 4) regardless of the relative strengths of their visual responses (bottom row).

We confirmed this observation statistically with permutation tests, conducted on pairwise latency differences separately for each contrast level and for each monkey (Methods). In monkey M, there were significantly longer latencies for bright stimuli in all contrasts (100% and 50% contrasts: mean differences = 3.18 ms and 3.56 ms, respectively, Monte Carlo p-values < 0.0001; 20% contrast: mean difference = 1.62 ms, Monte Carlo p-value = 0.0448; 10% contrast: mean difference = 2.03 ms, Monte Carlo p-value = 0.0152; and 5% contrast: mean difference = 4.5 ms, Monte Carlo p-value < 0.001). In monkey A, latencies were significantly longer for bright stimuli in 100%, 50%, and 5% contrasts (mean differences = 2.97 ms, 4.31 ms, and 4.91 ms, respectively; Monte Carlo p-values: < 0.0001, < 0.0001, and 0.0059, respectively); no significant differences were found in 20% and 10% contrasts (mean differences = 1.39 ms and 0.27 ms, respectively; Monte Carlo p-values: 0.0548 and 0.7587, respectively), but the same trends were still there (also see Fig. 5E). Thus, faster detection of dark than bright stimulus contrasts is a general property of SC neurons.

**Figure 5.**
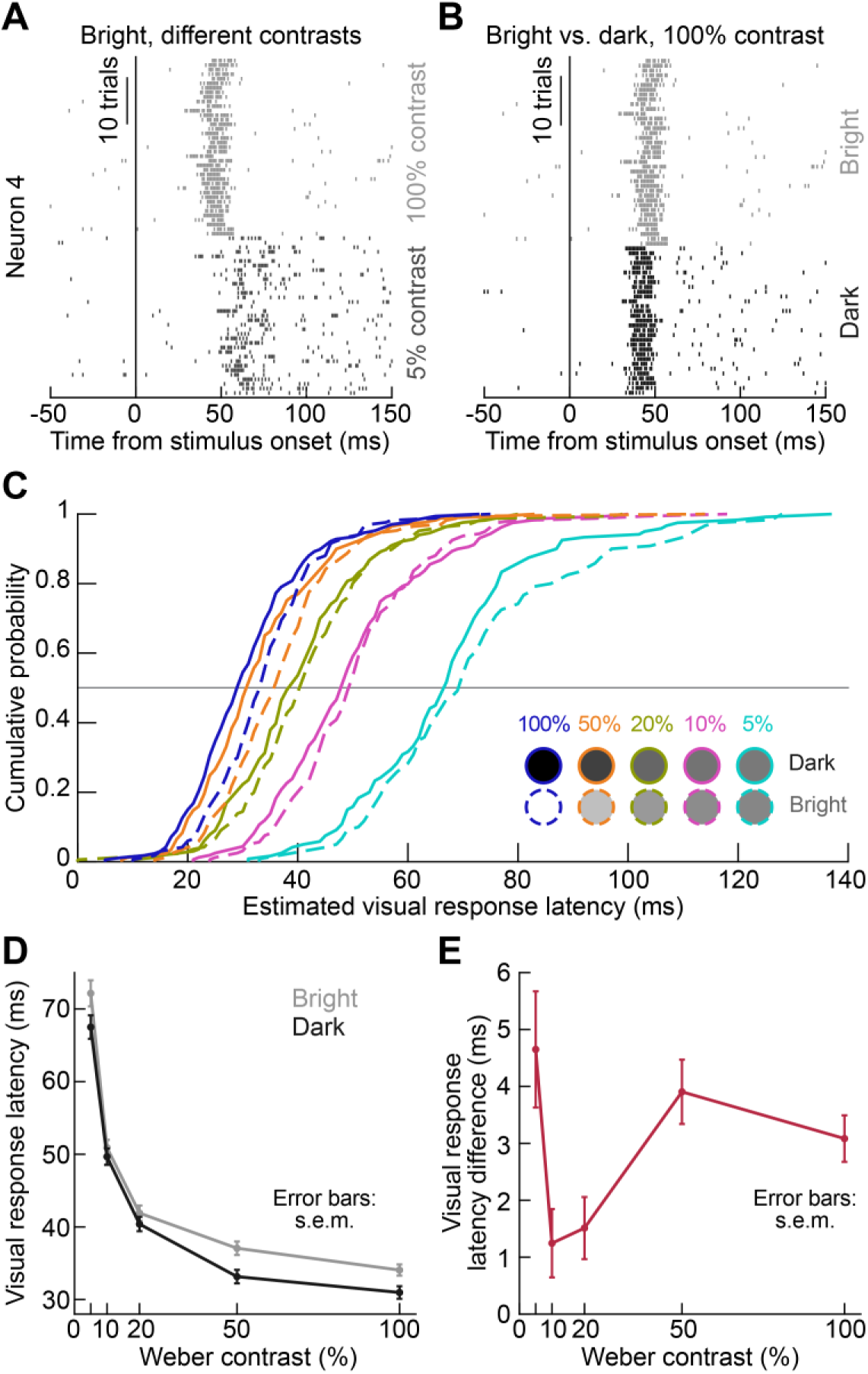
Interaction between contrast and luminance polarity in SC visual response latencies. **(A)** Visual responses of an example SC neuron from monkey M for high (100%) and low (5%) contrast bright stimuli (that is, with luminance higher than the background luminance). Each row of tick marks represents a single trial repetition, and each tick mark indicates the time of an action potential. The neuron responded earlier to the high contrast stimulus. **(B)** In the same neuron, the visual responses to a dark 100% contrast stimulus (that is, with luminance darker than the background luminance) occurred even earlier than the responses to a bright 100% contrast stimulus. Thus, both contrast and luminance polarity affected the neuron’s visual response latency. **(C)** Cumulative histograms of our estimates of visual response latency across all of our neurons, for both stimulus contrast (different colors) and stimulus luminance contrast polarity (solid versus dashed lines). High contrasts were associated with earlier visual response latencies in the SC. In addition, at each contrast, dark stimuli were systematically associated with earlier visual response latencies. (**D**) Average visual response latencies for brights and darks across contrast levels, demonstrating consistently faster responses for dark stimuli. (**E**) Average differences in visual response latencies between responses for bright and dark stimuli per contrast level. All contrast levels were associated with faster detection of dark than bright stimuli. The effect increased in strength with increasing contrast from 10% to 100%. At the 5% contrast condition, the effect was the strongest, likely because there were more dark-preferring neurons than at higher contrasts (see the bottom row of Fig. 4 and Fig. 6K, P). Error bars in **D**, **E** denote s.e.m. across neurons.

Of course, our results do not deny that high visual response sensitivity is normally associated with short visual response latencies. For example, with the sorting of neurons shown in Fig. 4 based on their response latency differences (top row), there was still a hint of an additional trend: neurons with a smaller latency difference between dark and bright stimuli tended to be the neurons preferring bright stimuli (bottom row). For example, compare the first and last quartiles in the bottom panel of Fig. 4A: more bright-preferring neurons occurred in the first quartile (having 0 or negative latency differences) than in the last quartile (having positive latency differences). This suggests that there were divergent forces influencing visual response latency: a neuron strongly preferring bright stimuli might have had its high response strength for bright stimuli (at a given contrast level) counterbalance the normally earlier detection of dark stimuli. Indeed, when we evaluated response latency as a function of both stimulus contrast (a proxy for visual response strength in the neurons) and luminance polarity, we found that both factors clearly influenced the neurons’ visual response latencies. This is shown in Fig. 5A, B for an example neuron, and in Fig. 5C for the population. In Fig. 5A, high contrast stimuli evoked stronger and, therefore, earlier visual responses than low contrast stimuli, as expected (Boehnke and Munoz, 2008; Marino et al., 2012; Marino et al., 2015; Hafed and Chen, 2016; Chen et al., 2018). With high contrast dark stimuli, the same neuron exhibited even earlier visual bursts than for high contrast bright stimuli (Fig. 5B). Across the population, cumulative histograms of estimated visual response latencies (Fig. 5C), as well as their averages and standard errors of the mean (Fig. 5D), revealed that increasing stimulus contrast systematically decreased response latencies, as expected (Boehnke and Munoz, 2008; Marino et al., 2012; Marino et al., 2015; Chen et al., 2018), but also that visual response latencies were always systematically shorter, at a given contrast, for dark than bright stimuli (consistent with Fig. 4; also see Fig. 6 below). This polarity effect on response latencies had an order of magnitude of a few milliseconds difference between dark and bright response latencies (Fig. 5E), similar to results in the cat LGN (Jin et al., 2011) and V1 (Komban et al., 2014). Therefore, both stimulus contrast (a proxy for response sensitivity) and stimulus polarity (conferring a temporal advantage for darks) dictated our SC neurons’ visual response latencies.

**Figure 6.**
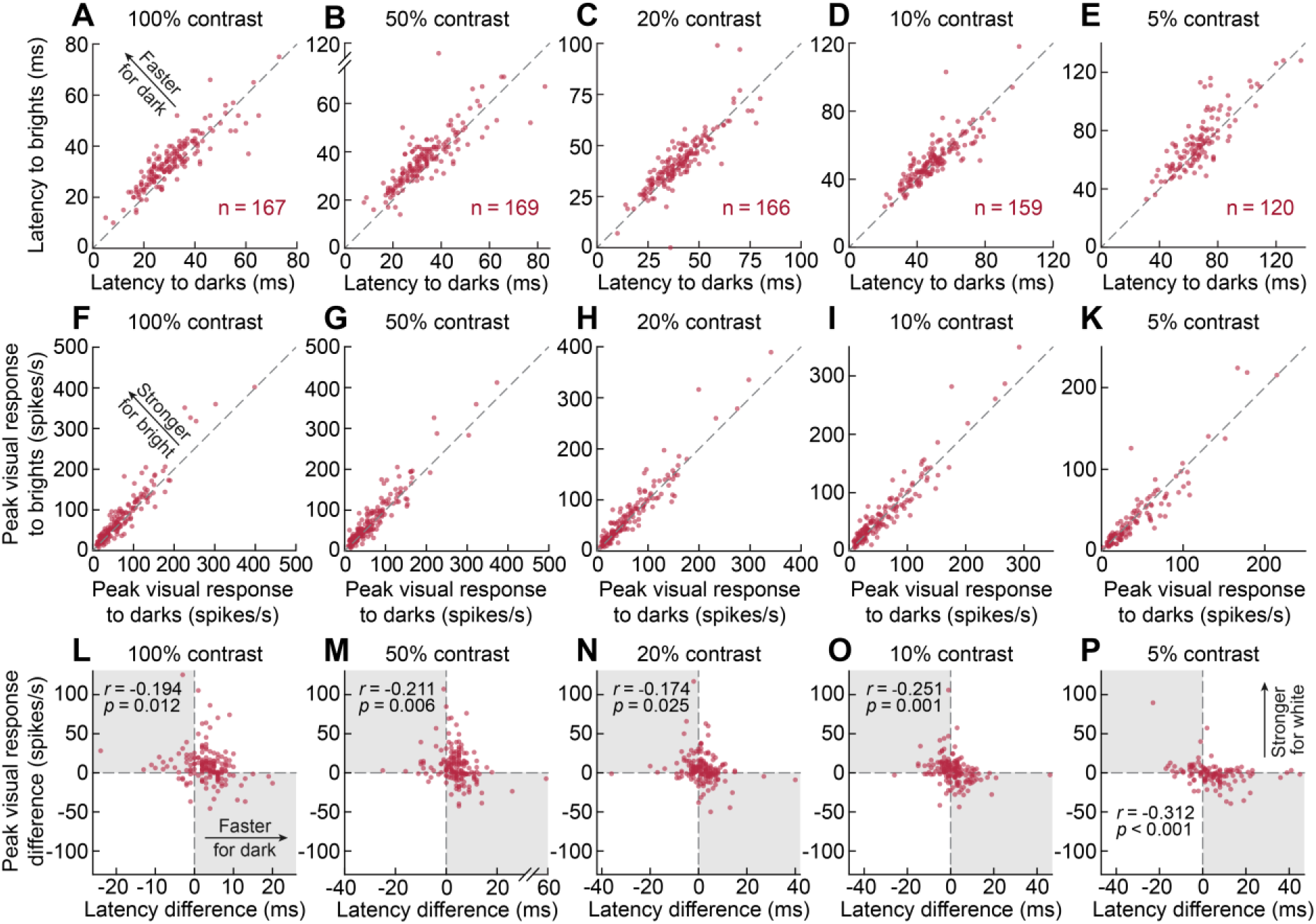
Contrast-dependent SC processing of dark and bright stimuli. **(A-E)** Scatterplots of response latencies to dark versus bright stimuli at each contrast level. The proportion of neurons responding faster to darks (above the unity line) increased with increasing contrast level. The proportion of neurons responding faster to darks was also substantially larger at the lowest contrast level (**E**). (**F-K**) Scatterplots of peak visual responses to darks versus brights at each contrast level. With increasing contrast, the proportion of neurons responding stronger to brights (above the unity line) also increased. (**L-P**) Correlations between differences in visual response latencies and differences in peak visual responses for brights and darks. The differences were obtained in the same way as in Fig. 4: a positive response latency difference indicates a faster response to dark stimuli, and a positive peak response means higher sensitivity to bright stimuli. The gray quadrants indicate the regions where preferences in terms of sensitivity and latency coincided: neurons in the upper left quadrant responded both stronger and faster for brights, and neurons in the lower right quadrant responded both stronger and faster for darks. There were weak negative correlations between response latency and sensitivity differences across contrasts, consistent with a known relationship between response latency and sensitivity. Critically, however, earlier responses to darks were not dictated by sensitivity preferences; if this was the case, most neurons would have occupied the shaded gray quadrants. Instead, at the highest contrast levels (e.g. **L**, **M**, **N**), the majority of neurons, which responded earlier for darks, responded stronger for brights (occupying the upper right quadrants), suggesting that dark stimuli were potent in expediting visual bursts despite the bursts being non-preferred by the neurons.

### Interaction between stimulus contrast and the processing of darks and brights by SC neurons

The results of Fig. 5E were particularly intriguing to us, in the sense that the lowest contrast stimuli (5%) were associated with a seemingly bigger effect of visual response latency difference between darks and brights than the more visible 10% and 20% contrast targets. One possibility could be that at 5% contrast, there were fewer bright-preferring neurons. We, therefore, next asked whether there was an interaction between luminance contrast level and the preference of neurons for darks or brights. To do so, we replotted the data above as scatter plots of visual response latency for brights versus darks in Fig. 6A-E and as scatter plots of visual response sensitivity for brights versus darks in Fig. 6F-K. The latency plots confirmed our earlier observations that there was systematically faster detection of dark contrasts across all contrasts. For sensitivity, there was an interaction between contrast level and SC population preference. At high contrasts (e.g. Fig. 6F, G), the population was biased toward preferring bright stimuli (despite responding faster for dark stimuli), whereas at 5% contrast (Fig. 6K), this bias disappeared and tended to be in the opposite direction; neuron 1 in Figs. 1D, 3A also demonstrates this effect: even though the neuron preferred brights in its plateau firing rate of the contrast sensitivity curve, its (weak) response at 5% contrast was still higher for dark targets. Thus, there was an interaction between contrast level and sensitivity to darks in our SC neurons. While this pattern is different from natural image statistics (Cooper and Norcia, 2015) and cat V1 properties (Liu and Yao, 2014), in the sense that we found more bright-preferring than dark-preferring neurons at high contrast, it does suggest that the larger latency effect magnitude in Fig. 5E at 5% contrast might have been driven by a larger number of dark-preferring neurons at this contrast level.

Statistically, we confirmed that there were contrast-dependent sensitivity preference differences between brights and darks. We applied the same pairwise latency difference procedure described above, but now to pairwise peak visual response differences (Methods). In monkey M, SC visual responses to brights were significantly stronger than responses to darks in the 100%, 50%, and 20% contrast conditions (mean differences = 11.69 spikes/s, 8.92 spikes/s, and 4.26 spikes/s, respectively; Monte Carlo p-values: < 0.0001, < 0.001, and 0.0187, respectively); the differences were not significant for 10% and 5% contrasts, and their trends were in the opposite direction (mean differences = −0.09 spikes/s and −2.35 spikes/s, respectively; Monte Carlo p-values: 0.9611 and 0.1712, respectively). In monkey A, the neurons were significantly more sensitive for brights in all but the lowest contrast (100%, 50%, 20%, 10%, and 5% contrasts: mean differences = 6.73 spikes/s, 7.35 spikes/s, 6.52 spikes/s, 6.92 spikes/s, and 0.81 spikes/s, respectively; Monte Carlo p-values: 0.0106, 0.002, 0.0028, < 0.001, and 0.8652 respectively).

In the same vein, for each neuron, we plotted the visual response latency difference against the peak response difference in Fig. 6L-P. We used the same conventions as in Fig. 4: a positive response latency difference indicating a faster response to dark stimuli, and a positive peak response difference meaning higher sensitivity to bright stimuli. If the results of Figs. 4, 5 were solely determined by response sensitivity at each contrast level, then all neurons should have occupied the shaded quadrants of these plots. In contrast, only a minority of neurons occupied these quadrants, particularly at high contrast. For example, in Fig. 6L, even neurons with >50 spikes/s difference in peak sensitivity in favor of bright stimuli were still significantly faster to detect dark stimuli. Interestingly, at 5% contrast, there were significantly fewer bright-preferring neurons, again providing a plausible explanation for the relatively large effect size in response latency seen in Fig. 5E at 5% contrast versus 10% and 20% contrast.

### Interaction between visual field location and the faster detection of darks by SC neurons

The above results indicate that there is faster detection of dark than bright stimuli by SC neurons, in general. However, it is also known that SC visual responses preferentially process the upper visual field (Hafed and Chen, 2016), consistent with the notion that eye movements support orienting towards or away from extra-personal stimuli largely occupying the upper visual field (Previc, 1990). If that is indeed the case, then it might be expected that differential temporal processing of dark versus bright stimuli might be magnified in the SC’s upper visual field representation. For example, birds of prey, or other threats, across a daylight sky would normally cast dark contrasts on retinal images, and they need to be detected efficiently by SC neurons. We, therefore, also asked whether the results of Fig. 4 could depend on the visual field locations of our recorded neurons.

We repeated the analyses of Fig. 4, but this time after separating neurons based on upper and lower visual field RF locations. The results are shown in Fig. 7 (for the highest contrast stimuli only, for simplicity). There was indeed a larger latency difference between dark and bright stimulus responses in the upper visual field neurons than in the lower visual field neurons (top panel). That is, the latency advantage for dark stimuli was magnified in the case of upper visual field SC neurons. We tested this observation statistically by using a permutation test with 10000 shuffles. Here we should note that although we pooled the data of both monkeys for visualization purposes in Fig. 7, we performed the statistical procedures only on data collected from monkey A. This was because there was a strongly unbalanced sampling of neurons in the upper and lower visual fields in monkey M (68 and 21 upper and lower visual field neurons in this monkey, versus a more balanced distribution of 24 and 33 neurons in monkey A). In monkey A, there was a significant difference between upper and lower visual field effects (mean difference = −4.09 ms, Monte Carlo p-value = 0.0099, Methods). Therefore, a known visual response latency advantage for the upper visual field in SC neurons (Hafed and Chen, 2016) was accompanied, at least in one monkey, by a larger difference between dark and bright stimulus responses. This result is consistent with the ecological likelihood of dark contrasts in natural environments (Liu and Yao, 2014; Cooper and Norcia, 2015), and also with the role of the SC’s visual processing machinery in supporting the sampling of extra-personal visual space by orienting eye movements (Previc, 1990; Hafed and Chen, 2016; Fracasso et al., 2022).

**Figure 7.**
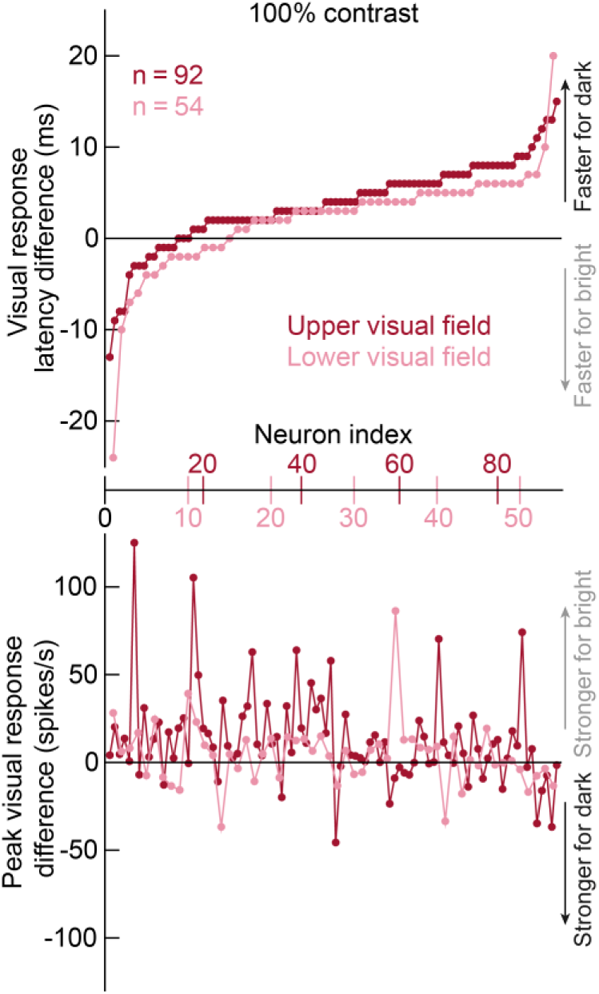
Magnification of response latency and sensitivity differences between dark and bright stimuli in the upper visual field. We repeated the same analyses of Fig. 4A, but now separating the neurons according to whether they represented upper or lower visual field locations. The top panel shows that the faster detection of dark stimuli by SC neurons that we saw in Fig. 4 was amplified for upper visual field neurons. The lower panel again shows that the effects of the top panel were dissociated from visual response sensitivity (all figure conventions are similar to Fig. 4). Interestingly, the lower panel shows that the absolute value of visual response sensitivity difference (between brights and darks) was also higher for upper visual field neurons than for lower visual field neurons (compare the dynamic ranges of the two curves). Therefore, both visual response latency and visual response sensitivity effects, in terms of luminance contrast polarity, were magnified in the upper visual field.

Note also that the same dissociation between response latency and response sensitivity occurred in Fig. 7 as in Fig. 4: the bottom panel in Fig. 7 shows that with the same ordering of the neurons as in the top panel, response sensitivity was not systematically ordered in either the upper or lower visual fields, consistent with the results of Fig. 4. Interestingly, the absolute value of the difference in response strength between dark and bright stimuli was also higher in the upper visual field neurons than in the lower visual field neurons (monkey A; mean difference = −13.3 spikes/s; Monte Carlo p-value = 0.0166, permutation test). This suggests that both latency differences (top panel) and absolute values of sensitivity differences (bottom panel) between dark and bright stimuli were amplified in the upper visual field representation of the SC, adding to a growing body of evidence of visual field asymmetries in the primate SC (Hafed and Chen, 2016; Hafed, 2021; Fracasso et al., 2022).

### Independence of the faster SC detection of darks from the luminance polarity at fixation

Finally, we wondered whether the black fixation spot at display center (Methods) might have dictated our results above. Such an effect would be unlikely because our neurons were extra-foveal; our stimuli were, therefore, generally far from the fixation spot (Methods). However, to unambiguously rule such an effect out, we repeated the same experiment with two slight modifications. First, the fixation spot was now white instead of black (Methods). If the black fixation spot was the reason for the faster detection of dark stimuli in the results of Figs. 1, 3–7 above, then this effect should be altered with a white fixation spot. Second, the fixation spot was now removed at the same time as stimulus onset, allowing the monkeys to generate immediate, visually-guided saccades. We analyzed 213 neurons recorded with this variant of the task (81 were also recorded in the original fixation task). We will describe saccadic reaction times as a function of contrast and luminance polarity in more detail below. However, for now, our aim was to replicate the visual burst results shown above. For each neuron, we normalized the neuron’s average firing rate by the peak visual response for 100% contrast stimuli in the interval 0-100 ms after stimulus onset (Methods). We then averaged all of the normalized firing rate curves of each monkey’s neurons (Fig. 8A, B). We separated the neurons of each monkey in this analysis to demonstrate the repeatability of our results across the animals, and also because subsequent saccadic behavior later in the trials differed between them, as we clarify in more detail below.

**Figure 8.**
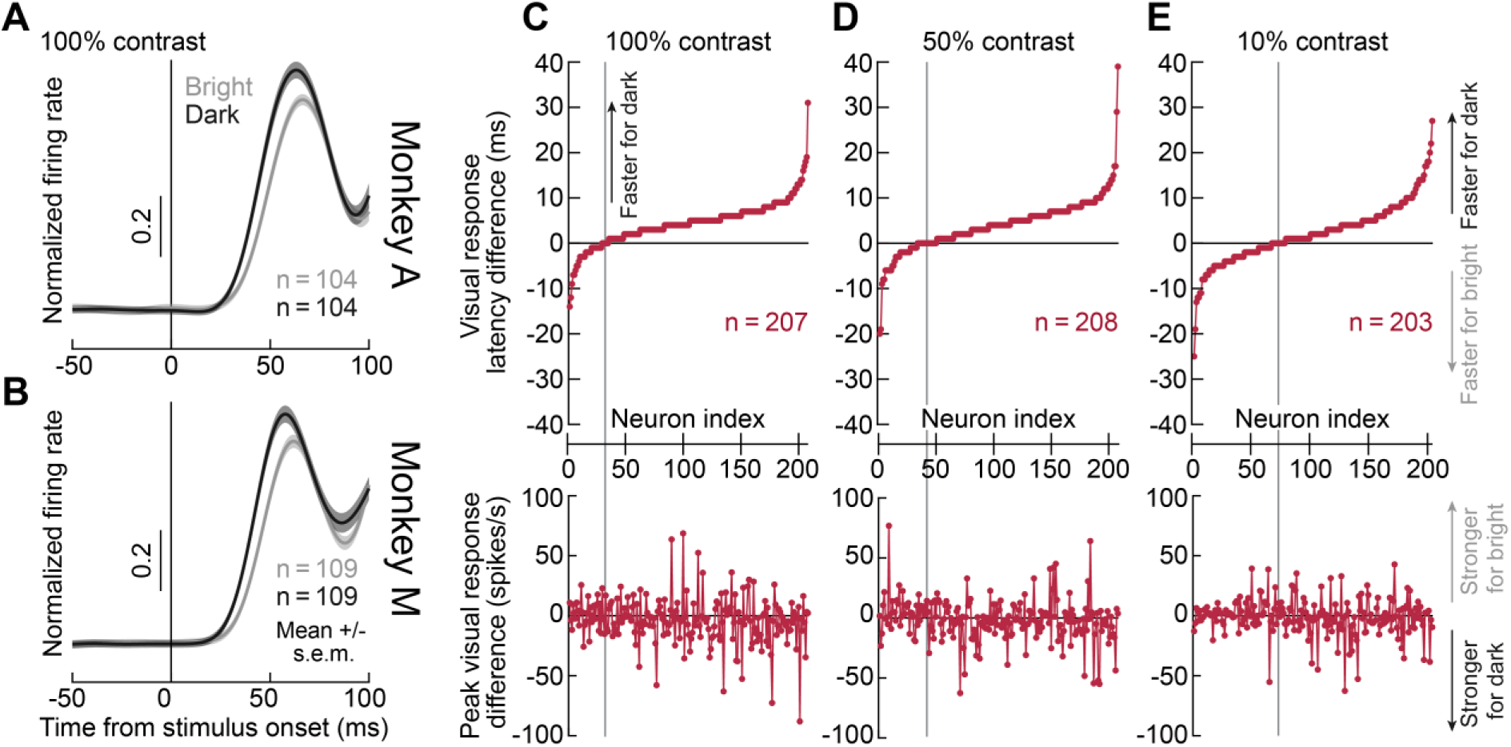
Faster detection of darks than brights in the SC, independently of central fixation spot appearance. **(A, B)** We repeated the same experiment as in Figs. 1–7, but this time with a white fixation spot rather than a black fixation spot (Methods). Also, here, we removed the fixation spot at stimulus onset, to allow monkeys to generate visually-guided saccades to the appearing stimuli. In each panel, we averaged each monkey’s stimulus-aligned firing rates (after normalizing each neuron’s firing rate curve to its peak visual response at high contrast; Methods). The numbers of neurons are shown in each panel, and error bars denote s.e.m. across neurons. As can be seen, visual responses still occurred earlier for dark stimuli than for bright stimuli in this task, and in both monkeys. Note that at around 100 ms from stimulus onset, there was an elevation of firing rates, which was the beginning of the saccade bursts for the triggered eye movements. Nonetheless, in monkey M, the elevation looked to be sharper than in monkey A, an observation that we discuss in more detail in Figs. 9, 11 below with respect to saccadic reaction times. **(C-E)** We replicated the analyses of Fig. 4 for the three contrasts that we tested in this task variant. The same conclusions were reached. Most neurons detected dark stimuli earlier than bright stimuli in their visual response latencies (top row), and this effect was dissociated from individual neuron sensitivity to either darks or brights (bottom row). All other conventions are similar to Fig. 4.

Both animals had clear visual responses in the task, consistent with the results of the fixation variant (Figs. 1, 3–7). Most importantly, these responses were also clearly still occurring earlier for dark stimuli than for bright stimuli (Fig. 8A, B). To summarize these results on an individual neuron basis, we replicated the same analyses of Fig. 4 (Fig. 8C-E; we combined the neurons of both monkeys here because of how similarly they behaved in Fig. 8A, B). We did this for all three stimulus contrast levels that we tested in this variant of the task (Methods). For all contrasts, most neurons still detected dark contrasts earlier than bright contrasts, irrespective of neural sensitivity (Fig. 8C-E), just as in Figs. 1, 3–7. Permutation tests run on the latency differences confirmed this observation for all contrasts (in monkey M, 100% and 50% contrasts: mean differences = 4.97 ms and 4.42 ms, respectively, Monte Carlo p-values < 0.0001; 10% contrast: mean difference = 1.62 ms, Monte Carlo p-value = 0.0185; in monkey A, 100% and 50% contrasts: mean differences = 3.71 ms and 3.35 ms, respectively; Monte Carlo p-values < 0.0001; 10% contrasts: mean difference = 2.52 ms; Monte Carlo p-value < 0.001). Note that the effect sizes were also of the same order of magnitude as those shown in Fig. 5E. Therefore, the results of Figs. 1–7 were not trivially caused by the use of a black fixation spot at screen center. Moreover, the results still persisted in a more reflexive behavioral task, in which prolonged fixation was not enforced in the face of a salient eccentric stimulus onset.

### Different temporal dynamics of firing rates in the sustained interval for darks and brights

The results so far have focused on initial visual bursts. However, with prolonged fixation (as in our primary task of Figs. 1–7), we also observed significant differences in SC neural response dynamics in the sustained interval (long after stimulus onset) for bright and dark stimuli. In particular, bright stimuli were generally associated with secondary elevations of firing rate above those of dark stimuli. To illustrate this, Fig. 9A, B shows the responses of four example neurons to high contrast stimuli (100%). Two neurons are from monkey A (Fig. 9A), and two neurons are from monkey M (Fig. 9B). In the first three neurons (neurons 5-7 in Fig. 9A, B), after the initial visual bursts, bright stimuli evoked stronger sustained activity than dark stimuli (see pink intervals highlighting the sustained interval). The stimuli were still present within the RF’s of the neurons in all cases, but there was an altered response dynamic after the initial visual bursts, particularly for bright stimuli. Even the fourth neuron (neuron 8 in Fig. 9B), which showed relatively weak sustained activity, still showed a subdued secondary peak in firing rate after the initial visual burst for bright stimuli (also see Fig. 11 below for more details on monkey M’s secondary bursts for bright stimuli).

**Figure 9.**
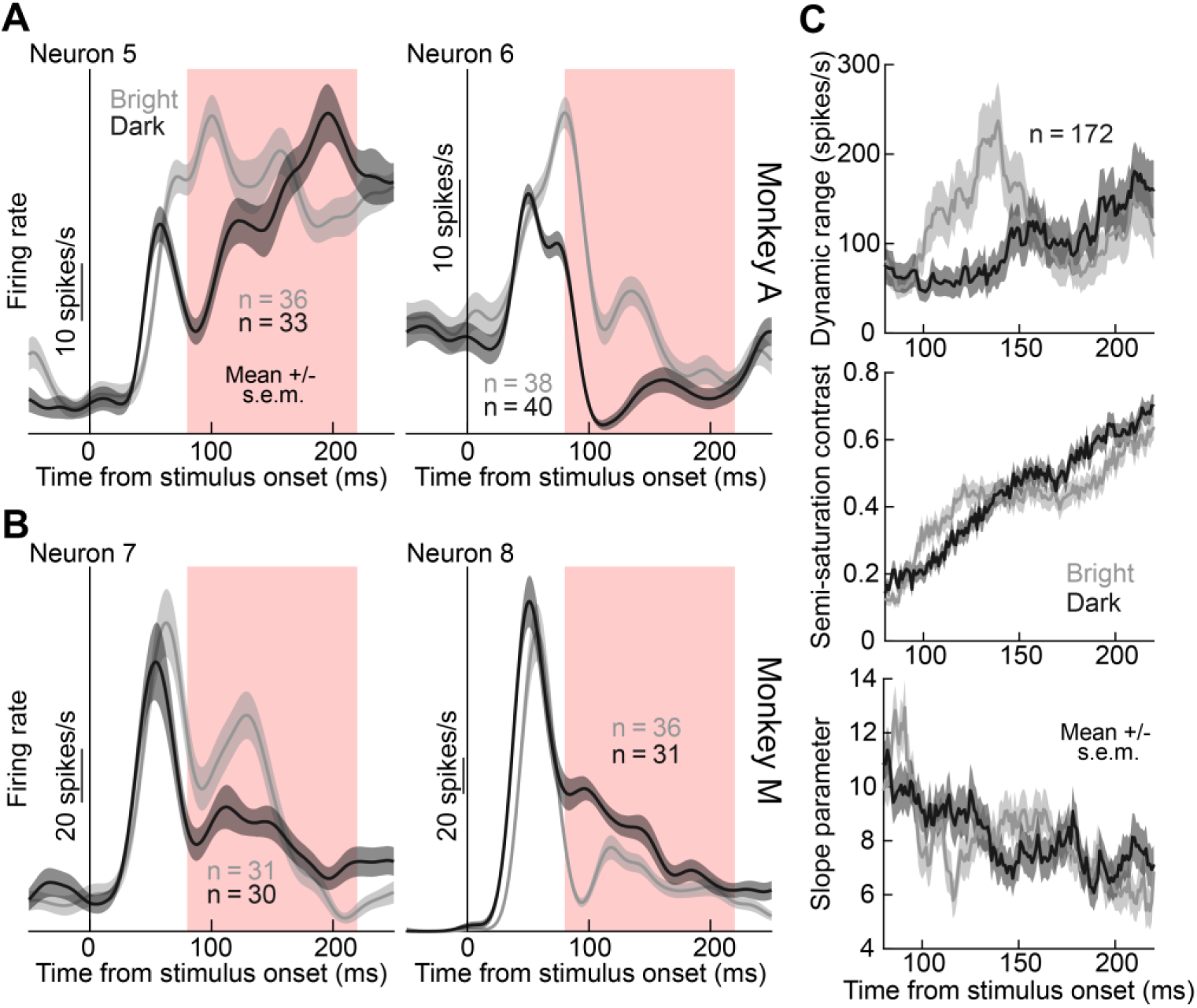
Preference for bright stimuli in later sustained intervals of visual neural SC responses. **(A)** Two example neurons from monkey A showing how later sustained responses (after the initial visual bursts) were elevated for bright more than dark stimuli. Responses for 100% contrast stimuli are shown, but similar observations were also made across contrasts (see **C**). **(B)** Two example neurons from monkey M showing similar observations. Note how neuron 8 had a weaker sustained response for bright stimuli, but it still exhibited a secondary burst (of small amplitude) for bright stimuli (also see Fig. 11D for this monkey’s population response summary in the sustained interval). The pink rectangles denote our interval of choice when analyzing sustained visual responses. **(C)** In such interval, we obtained millisecond-by-millisecond fits of contrast sensitivity curves for bright and dark stimuli. The dynamic range parameter of equation 1, *R*, showed a clear and significant elevation for bright stimuli relative to dark stimuli across the population (top panel). This was also the case in each monkey individually. The middle and lower panels show that the thresholds (middle panel) and slopes (lower panel) of contrast sensitivity curves were getting progressively worse in the sustained interval (relative to initial visual bursts), as expected, but with little differences between dark and bright stimuli. Error bars in all panels denote s.e.m. (across trials in **A**, **B**, and across neurons in **C**).

To characterize this altered dynamic of neural responses as a function of time in more detail, we took each firing rate curve after 80 ms from stimulus onset (that is, after the initial visual bursts). We then estimated contrast sensitivity curves at each time point. Each time sample of a firing rate curve is already a kind of average over some discrete measurement interval (due to the convolution of spike times with a gaussian kernel to generate firing rates). Therefore, we took each sample of the firing rate curve of a neuron in the sustained interval, and we used it to fit contrast sensitivity curves from equation 1 at each time point. This gave us a series of contrast sensitivity curves as a function of time. We then plotted the time courses of the parameters *R*, *C50*, and *N* of the fits during the sustained interval, and we did this for either bright or dark stimuli. The results across the entire population of neurons are shown in Fig. 9C. As can be seen, all parameters were varying differently between darks and brights in the interval around approximately 100-200 ms after stimulus onset (that is, during sustained visual response intervals), consistent with the example neurons of Fig. 9A, B. The biggest effect was in the *R* parameter, which was stronger for brights than darks, suggesting higher sustained firing rates for brights after the ends of the initial visual bursts. Both *C50* and *N* gradually changed in a manner that was consistent with higher thresholds and shallower contrast sensitivity functions in the sustained interval. That is, the contrast sensitivity of the neurons was generally the highest in the initial visual burst intervals, and it gradually degraded in sustained intervals (other than the *R* parameter elevation for bright stimuli). This makes sense given that sustained intervals were generally associated with much lower firing rates than in the initial visual burst intervals (and were therefore less likely to be strongly differentially modulated by stimulus properties). In any case, during the sustained interval, and unlike in the initial phases of SC visual responses, there was a generalized elevation of firing rates for bright stimuli compared to dark stimuli for all contrasts. As we show later, this effect was strong enough in monkey M, to the extent that it appeared to dominate this monkey’s saccadic reaction time patterns in the immediate, visually-guided saccade version of the task.

### Earlier detection of dark stimuli also by inhibited SC neurons

In all of the above analyses, we focused solely on neurons exhibiting positive visual responses (that is, increases in firing rates above baseline). However, with our offline neuron sorting pipelines (Methods), we also isolated a fewer number of neurons that exhibited transient decreases in activity after stimulus onset rather than increases. These neurons were obtained from similar recording sites to those from which we isolated neurons with visual bursts (we used linear electrode arrays primarily orthogonal to the SC surface; Methods). The neurons were, therefore, from similar topographic locations to those associated with the neurons reported in Figs. 1–9. When we analyzed these inhibited neurons in more detail, we found that their transient, stimulus-induced decreases in firing rates were still sensitive to luminance polarity. For example, in Fig. 10A, B, we show two example neurons from one of our electrode penetrations in monkey M. The two neurons showed classic visual responses (to black stimuli); moreover, their RF’s (shown in the insets for data collected with the presentation of black small spots during fixation) were spatially localized and overlapping with each other. From the very same electrode penetration, Fig. 10C shows a third sample neuron that was recorded simultaneously with the two other neurons; it was thus in the same SC topographic region as the two neurons of Fig. 10A, B. The neuron of Fig. 10C was inhibited instead. Most interestingly, this neuron clearly “responded” to a high contrast dark stimulus earlier than to a bright stimulus of the same contrast, with the only difference from the results of Figs. 1–9 being that the response in this case was a transient reduction from baseline activity rather than an increase. Across the population of such inhibited neurons (n=15 neurons), we repeated the same latency analyses of Fig. 4 above. That is, we assessed the relative time of “response” between bright and dark contrasts (Methods). As can be seen from Fig. 10D, the majority of such inhibited neurons also reacted to dark stimuli earlier than to bright stimuli, just like with the neurons possessing visual bursts. Similar observations were also made for the lower contrast stimuli. Interestingly, all 15 inhibited neurons had their “response” to stimulus onset slightly later than classic visual bursts in other neurons (compare the visual bursts in Fig. 10A, B to the inhibition time in Fig. 10C; the inhibition occurred slightly later than the bursts). Therefore, even inhibited neurons in the SC detected dark contrasts faster than bright contrasts.

**Figure 10.**
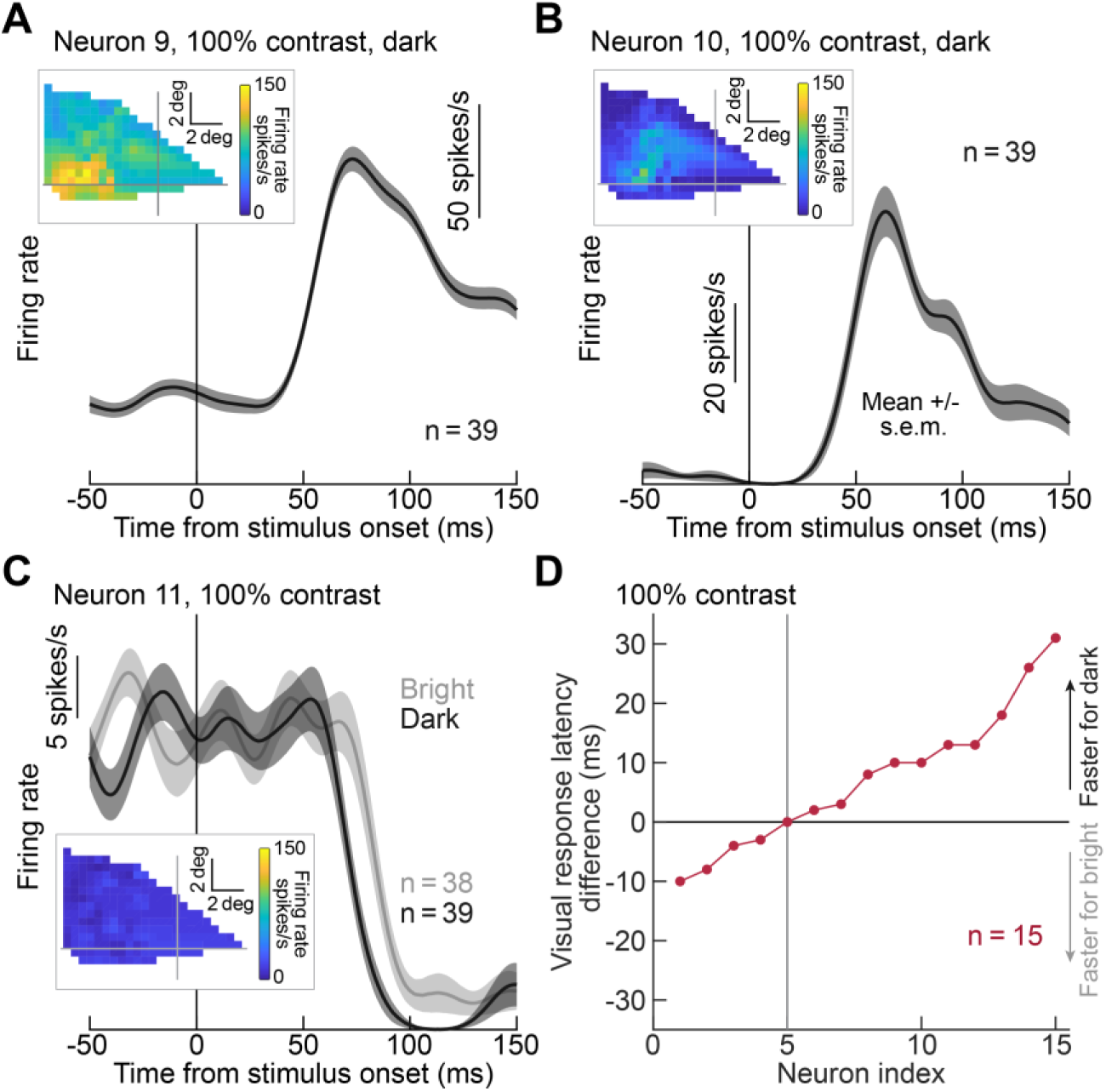
Faster detection of darks than brights even in SC neurons inhibited by stimulus onset. **(A, B)** Firing rates of two example neurons from a single linear electrode array penetration into the SC of monkey M. Both neurons had visual responses to dark stimuli in the upper left quadrant, with their RF’s (obtained by presenting small black spots at different locations; insets) being well localized in space, consistent with the SC topographic representation (Robinson, 1972; Chen et al., 2019). **(C)** A third neuron recorded simultaneously with the neurons in **A**, **B**. This neuron was inhibited by stimulus onset (also see the RF map in the inset). Nonetheless, the inhibition was still stimulus-dependent: there was earlier inhibition for dark than bright stimuli, consistent with our earlier results (e.g. Figs. 1, 4–8). Error bars denote s.e.m. across trials in **A**-**C**. **(D)** Replication of the analysis of Fig. 4A (top) for all neurons that were inhibited by stimulus onset. Now, we estimated visual response latency by checking when the neural activity was significantly decreased from baseline. Most neurons were still modulated earlier by dark than bright stimuli, consistent with our earlier results for visual bursts (e.g. Figs. 1, 4–8). All other conventions are similar to Fig. 4A.

### Saccadic reaction times can be significantly shorter for dark stimuli

Finally, prior work has demonstrated a tight relationship between SC visual response properties and saccadic reaction times (Boehnke and Munoz, 2008; Marino et al., 2012; Marino et al., 2015; Hafed and Chen, 2016; Chen et al., 2018). Specifically, both visual response sensitivity (Boehnke and Munoz, 2008; Marino et al., 2012; Marino et al., 2015; Hafed and Chen, 2016; Chen et al., 2018) and visual response latency (Chen et al., 2018) can predict such reaction times. Therefore, given the faster response latencies of SC neurons for dark stimuli that we found, we wondered whether this effect was sufficient to be associated with faster saccadic reaction times to such stimuli (even when response sensitivity was, on average, similar for darks and brights, as shown in Fig. 2; or even slightly favoring brights at high contrasts, as shown in Fig. 6). We tested our two monkeys and a third one on the immediate visually-guided saccade task; we used the same sessions as in Fig. 8 for monkeys M and A, and we ran separate behavior-only sessions for monkey F. The monkeys simply generated a saccade as soon as the target appeared (the fixation spot also disappeared at target onset, as mentioned above for Fig. 8 and in Methods). We measured saccadic reaction times and plotted them as a function of stimulus contrast and stimulus luminance polarity.

All monkeys showed faster reaction times for higher contrast stimuli, as expected (Marino et al., 2012; Marino et al., 2015) (p < 3×10^−103^ in each monkey individually, Kruskal-Wallis test exploring the effect of contrast on reaction time, when collapsing across luminance polarities). Interestingly, two out of the three monkeys (A and F) also showed consistently faster reaction times for the darker stimuli, like with the SC visual bursts. These results are shown in Fig. 11A, C, E; monkeys A and F were faster to react to dark stimuli at all contrasts. Monkey M, on the other hand, had faster reaction times for the bright stimuli (Fig. 11C). All of these results (of luminance polarity effects on reaction times) were significant (p < 2.5×10^8^ in each monkey individually, Kruskal-Wallis test exploring the relationship between luminance polarity and reaction time, when collapsing across contrasts); the effect sizes (shown for each condition in Fig. 11A, C, E) were also substantial. In addition, the effect sizes in monkeys A and F were of the same order of magnitude as the effect sizes of the visual response latency differences between darks and brights seen in Fig. 5E.

**Figure 11.**
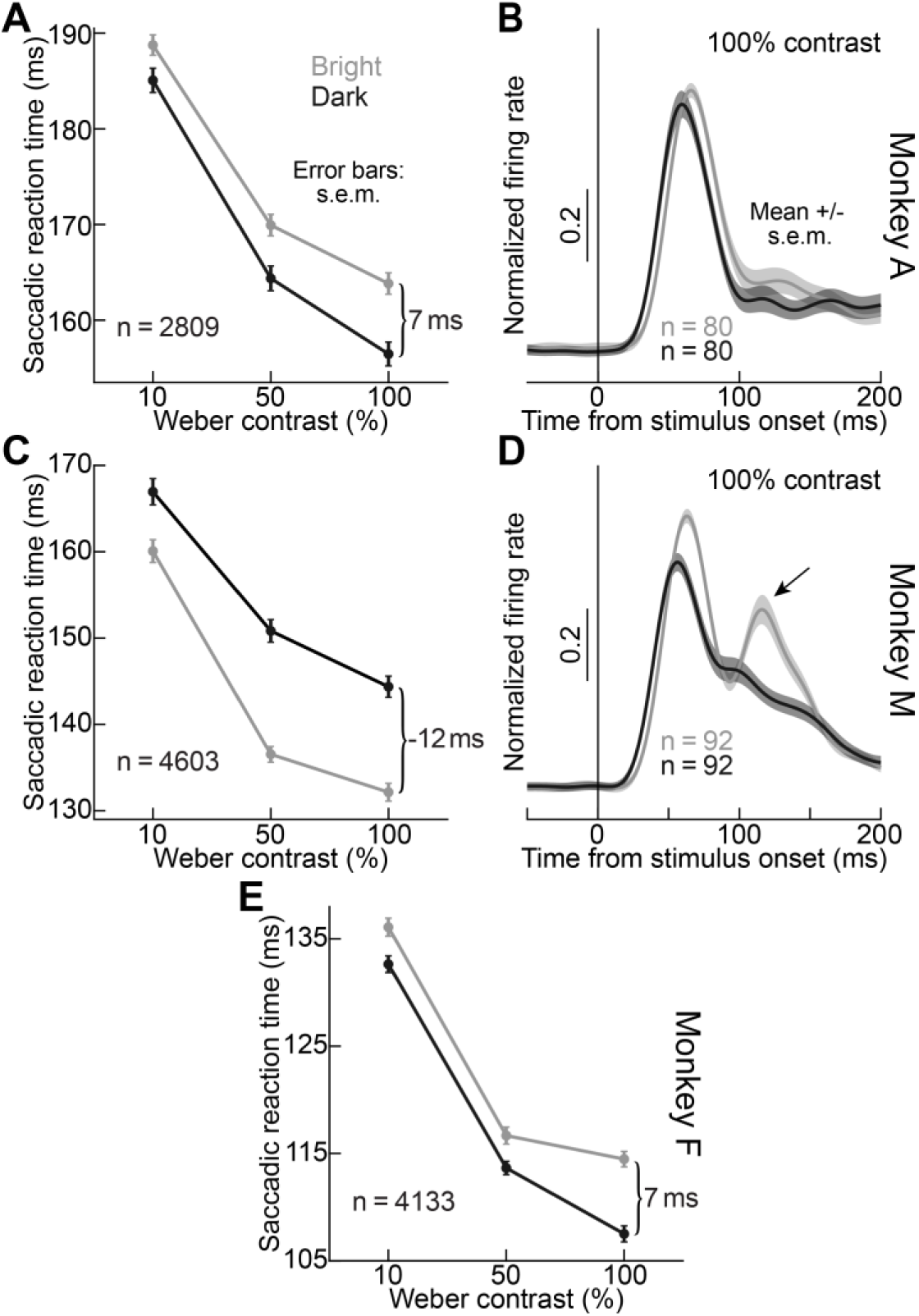
Relationship between saccadic reaction times and SC visual response properties for darks and brights in the SC. **(A)** Saccadic reaction times as a function of stimulus contrast (x-axis) and luminance polarity (different lines) in monkey A from the saccade version of our task. Error bars denote s.e.m. across trials. Reaction times were significantly shorter for dark than bright stimuli at all contrasts, suggesting a potential role for the earlier SC visual responses for dark stimuli in triggering earlier saccades for such stimuli. **(B)** This monkey’s neural responses during the fixation task showed clearly earlier visual bursts for dark stimuli, with equal visual burst strengths for darks and brights (consistent with Fig. 2). The figure was obtained similarly to Fig. 8A, B. Note also that the population sustained response (from the peak of the visual burst onward) was larger for brights than darks (consistent with Fig. 9). Error bars denote s.e.m. across neurons. **(C)** Monkey M showed the opposite reaction time effects from Monkey A. **(D)** In this monkey’s neurons, the secondary burst for bright stimuli in the fixation variant of the task was particularly prominent (compare to the monkey A neural responses). This suggests that in this monkey, this secondary preference for bright stimuli might have dominated the monkey’s reaction times in the saccade task. **(E)** We tested a third monkey behaviorally, and we replicated the monkey A results. Therefore, in 2 out of the 3 monkeys, saccadic reaction times were earlier for dark than bright stimuli, consistent with the neural results of Figs. 1, 4–8, 10.

We were particularly intrigued by the discrepancy in the reaction times of monkey M with respect to dark and bright stimuli. On the one hand, it might suggest that SC visual response latency (e.g. Figs. 4, 5) is not the only determinant of saccadic reaction times, which is indeed plausible. For example, we earlier found that SC visual response latency and visual response sensitivity together provided a better correlate of reaction times than either parameter alone (Chen et al., 2018). Therefore, since about half of the neurons in our population were more sensitive to bright stimuli anyway (Fig. 2), despite the faster detection of darks, it could be that this particular monkey’s reaction times were more dictated by SC visual response sensitivity than by visual response latency. On the other hand, it could additionally be the case that the later elevation of responses for bright stimuli that we saw in Fig. 9 was more pronounced in this monkey, potentially suggesting stronger top-down control for bright stimuli. In that case, bright stimuli could be preferentially processed by this monkey. Indeed, in a previous behavioral study in which we investigated the properties of saccadic inhibition as a function of luminance contrast polarity, this monkey reacted differently to full field white versus black visual flashes from the two other monkeys in the very initial oculomotor response to flash onset, again reacting faster for bright than dark flashes (Malevich et al., 2021) (see their Fig. 3). Therefore, we decided to check how this monkey’s neurons, in particular, reacted to white stimuli long after the initial visual bursts, and we were able to do so from our fixation variant of the task.

We plotted each monkey’s population visual responses for dark and bright stimuli in the fixation variant of the task, allowing us to explore the longer sustained interval. These results are shown in Fig. 11B, D. Even though monkey M’s neurons still detected dark stimuli earlier than bright stimuli in the initial visual response period (consistent with all of our results shown earlier), this monkey’s elevation of sustained visual activity for the bright stimuli (e.g. Fig. 9) was particularly pronounced when compared to monkey A (note the secondary peak in population firing rate for bright stimuli in Fig. 11D for monkey M, which was stronger than the same peak in monkey A). We also even saw hints of this secondary elevation in Fig. 8B in the immediate, visually-guided saccade variant of the task, with a sharper elevation for bright stimuli right after the initial visual burst and leading up to the saccade-related burst; however, of course, in this task, this sharper elevation for brights was harder to properly analyze in the saccade task because of how quickly the motor burst came.

Therefore, the results of Fig. 11 suggest that saccadic reaction times can indeed be faster for dark than bright stimuli, consistent with the faster detection of dark stimuli by SC neurons, and that even violations of such an observation (as in the case of monkey M) are still related to the SC visual responses (in this case, the sustained responses after the initial visual bursts subside).

In all, our results in this study indicate that SC neurons robustly detect dark stimuli faster than bright stimuli; that sustained visual responses in the SC instead favor bright stimuli; and that saccadic reaction times can reflect the faster detection of dark stimuli in the SC’s initial visual bursts and/or the later elevation for bright stimuli.

## Discussion

We evaluated the sensitivity of monkey SC neurons to luminance contrast polarity. We found that there was a diversity of preferences for darks and brights across the population (Figs. 1, 2). However, irrespective of preference (as defined by visual neural sensitivity), most neurons detected dark contrasts earlier than bright contrasts (Figs. 3–8, 10). Such earlier detection of dark stimuli was correlated with faster reaction times for such stimuli in 2 out of 3 monkeys (Fig. 11). And, even in the third monkey, this monkey’s observed opposite reaction time effect could be related to sustained modulations of SC neural activity, which exhibited a strong secondary elevation particularly for bright stimuli after the initial visual bursts (Figs. 9, 11).

Our results demonstrate that the primate SC does not necessarily exhibit identical ON/OFF sensitivity asymmetries for brights and darks as LGN and V1, refuting the idea that the primate SC simply inherits its visual properties from V1. For example, V1 neurons mostly prefer dark contrasts (Yeh et al., 2009), unlike in our SC population, and it would be interesting to further investigate whether deep V1 layers, projecting to the SC, violate this property or not. In fact, at high contrasts, our SC neurons significantly preferred bright, rather than dark, stimuli even while having faster response latencies to the dark ones (Fig. 6A, F, L).

Our results are additionally interesting because they add to a growing literature demonstrating that the primate SC is as visual a brain structure as the SC in other species, like mice, in which the SC is the primary recipient of retinal projections and, indeed, a primary visual structure. Consistent with this idea, the monkey SC receives a large amount of cortical visual input (Kadoya et al., 1971; Fries, 1984; Lui et al., 1995; Lock et al., 2003; Cerkevich et al., 2014), in addition to direct retinal input (Perry and Cowey, 1984). Thus, the SC in primates should be viewed as being even more visual than, say, the mouse SC. Such a rich visual nature of the primate SC matters a great deal for orienting responses, consistent with how SC visual responses can be linked to various aspects of saccadic behavior, like reaction time (Boehnke and Munoz, 2008; Marino et al., 2012; Marino et al., 2015; Hafed and Chen, 2016; Chen et al., 2018) and landing accuracy (Hafed and Chen, 2016). Such a link to saccadic behaviors was also clearly still evident in our current study (e.g. Fig. 11). In the future, it would be important to relate trial-to-trial variability in saccadic reaction times to trial-to-trial variability in SC, LGN, and V1 visual responses, as was done previously in V1 (Lee et al., 2010), to better appreciate the different functional specializations that exist in early visual responses that occur in multiple brain areas at approximately the same time.

Our observation that the primate SC can represent dark contrasts well (e.g. Fig. 2) is also consistent with earlier observations that SC neurons detect dark “shadows” (Humphrey, 1968; Cynader and Berman, 1972; Updyke, 1974). We are additionally particularly intrigued by the earlier response latencies for dark stimuli that we observed (e.g. Fig. 4), as well as by the altered temporal dynamics of responses during the sustained interval long after stimulus onsets (e.g. Fig. 9). These observations could potentially be used to further interpret earlier reports in the literature about SC visual and visual-motor modulations. For example, in investigating color-related responses in the SC, White and colleagues used a high contrast black target as the comparison stimulus to the colored ones (White et al., 2009). Because of that black stimulus, we predict that the latency differences that these authors observed relative to colored targets were slightly amplified than what they would have observed had they used a white target as the reference non-colored stimulus. Similarly, Churan and colleagues investigated how SC visual RF’s were modified around the time of saccades (Churan et al., 2011). They found dramatically different effects depending on whether the saccades were made across a gray background or across a dark background. It is intriguing to consider whether (and how) our sustained response effects, amplifying responses for bright stimuli (e.g. Fig. 9), could be related to their observations.

The fact that SC neurons can be strongly sensitive to dark contrasts is also interesting with respect to spatial frequency tuning in SC neurons. In recent work, we found that SC neurons can be sensitive to minute phase shifts of spatial frequency gratings, as small as 1 minute of arc in amplitude (Hafed et al., 2022). It would be fruitful, in light of these observations and the current work, to investigate RF subfield structure in more detail, for example, to study phase tuning in SC neurons. Indeed, both the current work and these recent results motivate a detailed mapping of RF’s with both bright and dark stimuli, to assess asymmetries beyond just visual response sensitivity and visual response latency. Indeed, prior work with reverse correlation techniques has suggested that there may be informative observations to be made about RF’s mapped with bright versus dark stimuli (Churan et al., 2012). In the near future, we hope to report SC RF maps for bright and dark stimuli in detail.

The altered long term temporal dynamics of firing rates as a function of luminance polarity that we observed (e.g. Fig. 9) also motivate modeling how these dynamics emerge. In V1, various stimulus factors, like contrast, alter not only the initial visual bursts (as might be expected), but also the sustained responses. Moreover, such alterations can be modeled using variants of linear/nonlinear filters and divisive normalization (Groen et al., 2021). It would be valuable to investigate such models in the SC, and to relate them to asymmetries in ON and OFF channels in both the LGN (Jin et al., 2011) and V1 (Komban et al., 2014).

Related to this, we would like to investigate, in the near future, how scene statistical regularities, with respect to orienting eye movements, can allow further expansion of our upper versus lower visual field analyses of Fig. 7. In these analyses, we were motivated by the theoretical framework of (Previc, 1990), in which he predicted that SC neurons should over-represent the upper visual field of retinal images because eye movements are relevant for sampling extra-foveal visual space. Thus, in the analyses of Fig. 7, we were driven by our earlier discoveries of significant asymmetries between upper and lower visual field SC neurons and saccadic performance (Hafed and Chen, 2016; Hafed and Goffart, 2020; Hafed, 2021; Fracasso et al., 2022). Indeed, we found that dark versus bright asymmetries were amplified in the upper visual field (Fig. 7), and this is ecologically sensible. For example, birds in the sky normally cast shadows on the retina. However, it would be even more intriguing to go even deeper when assessing such visual field anisotropies. For example, one could consider studying SC binocularity in more detail, to investigate whether neurons preferring far disparities would be more prevalent in the upper visual field representation of the SC or not. And, if so, would these far-preferring neurons also prefer more dark contrasts, like in the case of V1 (Samonds et al., 2012)? This is important to consider, especially given how our neurons seemed to prefer bright stimuli in a contrast-dependent manner (Fig. 6F-K) that was the opposite of what might be predicted from natural scene statistics (Cooper and Norcia, 2015).

Finally, we found a small subset of neurons, in the same topographic location as our bursting neurons, that were inhibited by stimulus onset (Fig. 10). Interestingly, these neurons still “detected” dark contrasts (by their transient inhibition of firing rate) earlier than bright contrasts. It would be important to investigate whether such neurons contribute to saccadic inhibition (Reingold and Stampe, 2002; Buonocore and McIntosh, 2008; Hafed and Ignashchenkova, 2013), which we recently found to also depend on stimulus luminance polarity (Malevich et al., 2021). Our initial intuition with regard to saccadic inhibition, described in detail in our theoretical proposal elsewhere (Hafed et al., 2021), is that structures beyond the SC are critical for this phenomenon. However, this does not deny the potential involvement of SC neurons (particularly those neurons that are transiently inhibited by stimulus onsets), and future research should investigate the mechanisms of saccadic inhibition in much more detail, including recording SC neurons with full or localized flashes of different luminance polarities like in psychophysics. Critical in those studies would be to quantitatively assess whether the small latency differences in “inhibition” versus excitatory visual “bursts” that we observed in Fig. 10 are consistent with the timing properties of saccadic inhibition or not.

## Acknowledgements

We were funded by the Deutsche Forschungsgemeinschaft (DFG): (1) BO5681/1-1; (2) EXC307; (3) HA6749/3-1; (4) HA6749/4-1; and (5) SFB 1233, Robust Vision: Inference Principles and Neural Mechanisms, TP 11, project number: 276693517.

